# Exploring the Modularity of *Thalassiosira oceanica* Aureochrome1 Photoreceptor-Transcription Factor: Characterization and Component Engineering

**DOI:** 10.1101/2025.03.07.642083

**Authors:** Anwesha Deb, Piu Sarkar, Devrani Mitra

## Abstract

The sensory transcription factors carry out gene expression in response to external stimuli like light. Further, they can be used in synthetic biological circuits to manipulate gene expression and thus can alter behavioral responses effectively. In this study, we characterize blue-light responsive photoreceptor transcription factor, Aureochrome (*To*Aureo1), from photosynthetic marine diatom *Thalassiosira oceanica*. *In silico* modeling and docking studies on C-terminal LOV (light-oxygen-voltage) and N-terminal bZIP (basic leucine zipper) domain (*To*AubZL) revealed conserved 3D structures as well as interaction with ‘Aureo-box’ containing substrate DNA. Modularity is an essential feature for synthetic biology inspired designs. Spectroscopy studies not only detected slow dark recovery kinetics, but also established the modular nature of both sensor and effector domains of *To*Aureo1. First, to check the compatibility of *To*Aureo1 bZIP with other non-Aureochrome LOV sensors; second, to explore the essentiality of heterogeneous linker connecting effector and sensor domains; and third, to generate temporal scaffolds for optogenetics, chimeric photoreceptor-transcription factor, *AsTo*bZL was generated. Upon blue-light illumination, both the wildtype and synthetic constructs showed a four-fold increase in DNA binding activity. Formation of higher order structures in both cases could be important for transcription activation. Their dark recovery time, however, varied significantly by about ten-fold – further establishing modularity in LOV sensors. Our design further proves the non-essentiality of native *To*Aureo1 linker between LOV and bZIP in blue-light mediated DNA binding, thereby opening up newer options in optogenetics. Compatibility among different *To*Aureo-bZIP partners further strengthens their possible role in light-regulated transcription factor interaction networks through large-scale participation.

## 1. Introduction

<u>T</u>ranscription factors (TFs) perform crucial roles in gene expression either by gene activation or silencing. Rational design of TF to enable need based DNA binding will not only facilitate gene therapy but will provide a useful tool for research in life sciences (Pomerantz et al., 1995). Transcriptional regulation can be driven by sensory TFs, which can be activated by diverse external stimuli e.g. light. Given the limited presence of light/photo-activated TFs in selective life forms, design of the same bears tremendous potential. The modularity of the photosensor domain, switching on/off gene expression with readily available and largely non-toxic light, makes the design of photo-activated TFs experimentally feasible. The reversible nature of dark-light photocycle in photoreceptors, automatically imparts ‘reversibility’ to the process, unlike those mediated by DNA/RNA caging or silencing through siRNA [(Dmochowski & Tang, 2007; Edwards et al., 2009; Gorostiza & Isacoff, 2008; Mayer & Heckel, 2006; Mikat & Heckel, 2007; Pinheiro et al., 2008; Shestopalov et al., 2007)]. Reversible photo-control can also be achieved by using azobenzene chromophore linked to DNA or modified peptides/proteins [(Caamaño et al., 2000; Dohno et al., 2009; Guerrero et al., 2005; Liang et al., 2002; Morgan et al., 2010)]. However, such a system is complex and is difficult for targeted therapy. The high diffusion rate of caged low molecular weight molecules may also unnecessarily trigger cross reactivity and activate some other molecule in the process (Möglich & Moffat, 2010). Therefore, naturally occurring light-activated TFs and synthetic TFs derived from them, seem to be better alternatives to achieve light-driven transcriptional regulation.

The photosensor module for designing synthetic light-activated TFs can be chosen from naturally occurring photoreceptors. Photoreceptors can be broadly classified into ten major groups, whose absorption wavelengths span across visible spectrum: Cryptochrome, Rhodopsin, Xanthopsin, Phytochrome, Cyanobacteriochrome, Car-H (adenosylcobalamine containing photoreceptor), OCP (<u>O</u>range <u>C</u>arotenoid <u>P</u>rotein), BLUF (<u>B</u>lue <u>L</u>ight <u>U</u>tilising <u>F</u>AD), UVR8 (<u>UV</u>-resistance Locus <u>8</u>, responding to UV-B range i.e 280-320 nm) and LOV (<u>L</u>ight <u>O</u>xygen <u>V</u>oltage) [(A. Banerjee, Herman, Kottke, et al., 2016a; Chaves et al., 2011; Christie et al., 1999; Losi & Gärtner, 2011; Rockwell & Lagarias, 2010)]. The prime advantage of genetically encoded photoreceptors is their specificity and *in situ* penetrability (Möglich & Moffat, 2010). Cells harboring exogenous genes encoding light-sensitive proteins (e.g. channelrhodopsins), can be specifically expressed by light, using optogenetics. With paramount potential to control multifarious cellular events with light at precise spatio-temporal resolution, optogenetics first emerged to address neuro-pathological events (Boyden et al., 2005). However, with the accommodation of multiple photoreceptors, in their natural and synthetic forms, optogenetics has now transcended the barrier of neuroscience. Recent years have even witnessed the advent of ‘plant optogenetics’ (Banerjee & Mitra, 2020), even though plant systems are difficult to engineer because of their inherent light-sensitivity. The production of plant biomass could be effectively enhanced through photoreceptor engineering and optogenetics [(Hart et al., 2019; Papanatsiou et al., 2019)]. With several advantages like endogenous protein production, high quantum efficiency as well as high dynamic range of flavin chromophore, flavin based photoreceptors are ideal candidates to design optogenetic scaffolds (Banerjee & Mitra, 2020; Möglich & Moffat, 2010) This study therefore focuses on flavin based LOV photoreceptor aureochrome, exclusively present in photosynthetic stramenopiles. Aureochromes (Aureos), first isolated and characterized from *Vaucheria frigida* are <u>b</u>asic leucine <u>zip</u>per (bZIP) containing LOV photoreceptor-TFs [(Mitra et al., 2012; Takahashi et al., 2007)]. Unlike most other LOV photoreceptors, Aureos possess an inverse domain topology - LOV sensor at the C-terminus and bZIP effector domain at the N-terminus. However, such effector-sensor domain architecture is similar to eukaryotic PAS (<u>P</u>er-<u>A</u>rnt-<u>S</u>im) containing TFs like HIF-1α, ARNT etc., crucial for the regulation of several genetic programs. LOV belongs to the PAS superfamily and bears the signature five-stranded antiparallel β-sheet in 2-1-5-2-3 topology with intermittent α-helices [(Crosson & Moffat, 2001; Wu et al., 2015)]. Sandwiched between β-core and α-helices, the flavin (FMN/FAD) chromophore interacts with cysteine *via* thio-ether bond formation upon blue-light illumination [(Möglich et al., 2009; Salomon et al., 2000; Taylor & Zhulin, 1999)]. Following withdrawal of light, LOV returns to the dark state. The time span of dark-reversal varies widely among different LOV photoreceptors, even within the same class like Aureos. This feature is in fact critical for the generation of temporal variants in optogenetics [(Deb et al., 2020; Hart et al., 2019)]. In this study, we first characterize <u>Au</u>reo1 from marine photosynthetic diatom *<u>T</u>halassiosira <u>o</u>ceanica* (*<u>To</u>*<u>Au</u>reo1), particularly the <u>bZ</u>IP + linker + <u>L</u>OV module (*To*AubZL), excluding N-terminal extension, using biochemical and biophysical characterization. We further check the compatibility of the Aureo effector (bZIP) with one most well employed LOV sensor in optogenetics and design a new chimeric photoreceptor-TF with different temporal control. Characterization, design and engineering of *To*AubZL, carried out simultaneously, allow us to explore the potential of Aureo-based optogenetic designs. It further helps us to gain deeper mechanistic insights into Aureo-driven TF interactions in *Thalassiosira* to promote light-activated gene expression.

## 2. Materials and methods

### 2.1 *In silico* characterization

#### 2.1.1 Sequence retrieval and analysis, Homology modeling and Docking

Aureochrome sequences from *Thalassiosira oceanica*, including *To*Aureo1, were retrieved from UniProtKB. We investigated the apparent disorderliness in *To*Aureo1’s big N-terminal extension (NTE) and its potential to serve as transactivation domain (TAD) using EXPASY ScanProsite as well as IUPred2A/AIUPRED server [(Erdős & Dosztányi, 2024; Mészáros et al., 2018)]. In absence of any experimental data, preliminary structural investigation of NTE was carried out using the available 3D structure from AlphaFold (Jumper et al., 2021) and secondary structure prediction software, PSSpred (Yan et al., 2013). Considering the unavailability of physiologically relevant dimer structure in AlphaFold database, we proceeded further with the *in silico* modeling of *To*Aureo1 LOV and bZIP domains separately using Discovery Studio and SWISS-MODEL software respectively. The full-length homology model of *To*AubZL could not be generated because of the absence of any information on the template structure against the linker region connecting bZIP and LOV. 1N9L (dark state LOV structure of *Chlamydomonas reinhardtii* Phototropin LOV1 domain) was taken as the template along with its non-covalently bound FMN, to model the *To*Aureo1 LOV fragment. The bZIP domain of *To*Aureo1 was modeled using 1DH3 (CREB TF) as the template. The model structures were validated through Ramachandran plot analysis in Molprobity (Williams et al., 2018). After generating two separate bZIP polypeptide chains, complementary bZIP strands were joined, following quaternary structural orientation of 1DH3, to get functional dimeric bZIP domain using *Pymol*. The energy minimization was conducted by *Chimera* (Pettersen et al., 2004), using default parameters i.e. 100 steepest descent steps followed by 10 conjugate gradient steps, where the step size is 0.02Å. The energy-minimized *To*Aureo1 bZIP was then docked with ‘Aureo-box: TGACGT’ containing CRE-substrate DNA from 1DH3 using *NPDock* software (Tuszynska et al., 2015) after pre-defining DNA binding basic region. After selecting the structure with the best docking score, the protein-DNA interactions were visualized and analyzed using *Pymol*. The flavin binding site of *To*Aureo1 LOV was further compared with other LOV counterparts from all Aureos e.g. *Vf*Aureo1 (Mitra et al., 2012), *Pt*Aureo1 (Heintz & Schlichting, 2016), *Od*Aureo1 (Kalvaitis et al., 2019) as well as *As*LOV2 (Halavaty & Moffat, 2007) using LigPlot+ (Laskowski & Swindells, 2011).

#### 2.1.2 Helical wheel projection and analysis of inter-bZIP interactions

To check the compatibility of *To*Aureo1 bZIP monomer with other bZIP monomeric units from *Thalassiosira* Aureos (as retrieved from UniProt KB), helical wheel projection models were developed. The models were developed manually after careful observation of the sequence alignment, conducted by *Clustal Omega*. Compatibility analysis is useful not only to predict intra-Aureo interactions but also to understand the essential presence of Aureos in photosynthetic heterokonts.

### 2.2 Cloning

#### 2.2.1 *To*Aureo1 LOV and *To*AubZL

The codon optimized gene sequence [corresponding to 276-520 amino acid (aa) residue stretch of K0SY93] of *Thalassiosira oceanica* Aureochrome1 (*To*Aureo1) was ordered from *GeneArt Strings* (*Invitrogen*). The following primers (*IDT*) were used for the amplification of *To*AubZL (bZIP + LOV domains; 276-520 of *To*Aureo1) and *To*Aureo1 LOV (LOV domain; 359-520): *To*AubZL_FP: 5’-GAT CGG ATC CAT GAG CAA CGC GGG C-3’; ToAubZL_RP: 5’-GTA CCC TAG GTT ACG CAT CCG CTT CAT C-3’; *To*AuL_FP: 5’-GGT CGC TAG CAT GCA GGC GCT GGC GG-3’;*To*AuL_RP: 5’-GTA CCC TAG GTT ACG CAT CCG CTT CAT C-3’;. Both the fragments (*To*Aureo1 LOV and *To*AubZL) were amplified by standard PCR protocols. Both the vector (pET28a) and inserts (*To*Aureo1 LOV and *To*AubZL) were double digested using NheI and SacI. The double digested products were cleaned and ligated with T4 DNA ligase (*New England Biolabs*) followed by transformation in *E. coli* DH5α cells. The purified plasmid DNA was verified by DNA sequencing (*Eurofins Genomics India Pvt. Ltd.*)

#### 2.2.2 *AsTo*bZL [*To*Aureo1 bZIP (corresponding to 303-351 aa sequence) + *Avena sativa* Phot1 LOV2, *As*LOV2 (corresponding to 404-546 aa sequence)]

For the construction of chimeric construct, the codon optimized LOV2 gene of *Avena sativa* phototropin1 was ordered from *GeneArt Strings* (*Invitrogen*). Both the fragments were amplified separately creating sticky overhangs at the point of fusion followed by a fusion PCR. The reaction was setup using the following primers: Forward primer of ToAureo1 bZIP: 5’-GAT CGC TAG CAT GCG CCG CGA ACG CAA C-3’; Reverse primer fusion overhang: 5’-GGT GGT CGC CAG CAT TTC TTT TTC GCC CAG-3’; Forward primer of fusion overhang: 5’-CTG GGC GAA AAA GAA ATG CTG GCG ACC ACC-3’; Reverse primer of *As*LOV2: 5’-GAT CGA GCT CTT ACA GTT CTT TCG CCG CTT C-3’. Both the fragments (corresponding to the above mentioned portions from *To*Aureo1 bZIP and *As*LOV2) were separately amplified by PCR. Both the fragments thus generated were then fused by PCR, which involved steps same as *As*LOV2. Both vector pET28a and fusion product (*AsTo*bZL) were double digested using NheI and SacI followed by ligation. The recombinant plasmid was transformed and extracted as in case of *To*AubZL.

### 2.3 Protein purification

Protein purification for all the three constructs [*To*Aureo1 LOV, *To*AubZL and *AsTo*bZL] followed the same methodology. Respective plasmid DNA was first transformed into *E. coli* C43 (DE3) competent cells and plated onto Kanamycin containing LB (Luria Bertani) plates. A single colony was picked up to inoculate 20 ml of starter culture (LB media with 50μg/ml kanamycin) and kept for overnight incubation at 37^0^C. The following day, 2 litres of LB media was inoculated using 1% starter culture and incubated at 37^0^C, 150 rpm till O.D._600nm_ reached ~0.6. The culture was next subjected to protein induction with 300 μM Isopropyl ß-D-1-thiogalactopyranoside (IPTG) (Promega/SRL). Induced culture was kept at 22^0^C for 16 hours. The bacterial cells were then harvested by centrifugation at 8000 rpm for 10 minutes. The cell pellet was either used immediately for purification or stored at −80^0^C for future use.

For protein purification, the harvested cells were resuspended in a lysis buffer (20 mM Tris, pH 8.0; 50 mM NaCl; 10% glycerol) followed by addition of lysozyme and Protease cocktail inhibitor (*Sigma*). The treated cell suspension was then sonicated and centrifuged at 12500 rpm for 1 hour. As all the constructs contained hexa-His tag, affinity chromatography was used as the first step of purification. The supernatant was first incubated with FMN and pre-equilibrated Ni-NTA resin (*Qiagen*) using batch binding method. The slurry was next passed through the empty column (*Thermo Scientific*) under gravity separation and flow-through was collected. Then the column was rinsed using a wash buffer (20 mM Tris, pH 8.0,; 50 mM NaCl; 10% glycerol; 10 mM imidazole). Freshly prepared elution buffer (20 mM Tris, pH 8.0; 50 mM NaCl; 10% glycerol; 250 mM imidazole) was then used to elute the desired protein fractions. The eluted fractions were concentrated using *Amicon* ultra filtration units (MWCO - 10KDa) to conduct either desalting or gel filtration chromatography. Desalting was done using the PD10 desalting column using the same equilibration buffer. For gel filtration chromatography, 50μl of concentrated protein was injected through ENrich gel filtration column (*Bio-rad*). The eluted fractions with desired protein samples were pooled for measurement of concentration (using absorbance at 447 nm) as well as dark recovery kinetics using UV-Vis spectroscopy. In other batches of protein purification (especially meant for EMSA studies), the desalted protein (*To*AubZL/*AsTo*bZL) was loaded onto HiTrap Heparin HP column (*Cytiva*) using the same equilibration buffer as gel filtration/desalting and eluted using a 50mM - 1.2M NaCl gradient. The desired protein fractions were desalted again and concentrated for further use.

### 2.4 UV-Visible spectroscopy and dark recovery kinetics

UV-Vis spectroscopy of *To*Aureo1 LOV, *To*AubZL and *AsTo*bZL was conducted for each sample to measure the concentration of proteins from different batches of purification as well as dark recovery kinetics. Calculation of protein concentration was carried out using OD_447nm_ in the dark state, considering the extinction coefficient of FMN [12500 M^-1^ cm^-1^]. The absorption maximum at 447 nm with flanking shoulders is typical of LOV photoreceptors. For dark recovery kinetics, absorption spectra for all three constructs were recorded by UV-Vis double beam spectrophotometer (*Shimadzu*, UV-2600) for the range 230-800 nm. The spectra were recorded both in the dark state and after 10 minutes illumination using 364 W/m^2^ LED light. Dark recovery kinetics was then conducted for each sample till no further recovery at 447 nm. The OD_447nm_ values were then plotted against time using a single exponential equation [OD_447nm_ = constant(1-e^-*kt*^] to calculate the rate constant (*k*) and lifetime of the constructs.

### 2.5 Circular Dichroism spectroscopy

*To*AubZL and *AsTo*bZL: 1.9 mg/ml and 0.88 mg/ml concentration of proteins were used for *To*AubZL and *AsTo*bZL respectively. Far-UV data (190-250nm) was collected in a CD spectropolarimeter (JASCO, J-815), using a 1mm Quartz cell. Data points were recorded using 1nm step resolution, a time constant of 2 seconds, a scan speed of 50nm/min and a spectral bandwidth of 2nm. The CD spectrum was analysed from DICHROWEB (Sreerama & Woody, 2000), using SELCON3 and CONTIN-LL algorithms.

### 2.6 Electrophoretic mobility shift assay

*To*AubZL and *AsTo*bZL: “TGACGT” or Aureo-box (A. Banerjee, Herman, Serif, et al., 2016; Takahashi et al., 2007) containing substrate DNA strands were ordered from IDT. Annealing buffer (20 mM Tris, pH 8; 30 mM NaCl; 10 nM substrate DNA-strand 1; 10 nM substrate DNA-strand 2) was used to anneal the single DNA strands after initial heating at 95^0^C. The mixture was left overnight for slow cooling and double-stranded substrate DNA was obtained. ‘Aureo-box’ (highlighted) containing DNA template 5’-TGT AGC GTC <u>TGA</u> <u>CGT</u> GGT TCC CAC-3’ and its complementary strand were used for both *To*AubZL and *AsTo*bZL. EMSA was set up using varied concentration of previously purified protein fractions in binding buffer (20 mM Tris, pH 8; 50 mM NaCl; 2 mM MgCl_2_; 20% glycerol) keeping DNA concentration (0.4 μM DNA) constant in all tubes. Concentration of proteins were adjusted by serial dilution. EMSA was performed under both light and dark conditions for *To*AubZL and *AsTo*bZL. The dark and light state EMSA were conducted separately under red-light or white-light conditions, to prevent any influence from the opposing states. Following incubation for 20 minutes, the samples were next loaded onto 10% native PAGE and resolved at 150 V for ~40 minutes. The gel was stained with SyBr gold DNA stain (*Thermo Scientific*) in 0.5X Tris-borate-EDTA buffer and the image was collected using Chemidoc (*Bio-rad*). The consistency in DNA binding pattern under dark and light conditions for both *To*AubZL and *AsTo*bZL was validated by performing EMSA with samples from multiple batches of protein purification.

## 3. Results

### 3.1 Sequence retrieval of multiple Aureochrome sequences from *Thalassiosira oceanica*

While searching for aureochrome gene from *Thalassiosira oceanica* (*To*Aureo), we came across multiple entries in the UniProtKB. We obtained the sequences and ran a multiple sequence alignment with other Aureo sequences **[Figure S1]**. Based on the similarity of sequences [**Figure S2**], we designate K0SY93 as *To*Aureo1, A0A6S9WQI9 as *To*Aureo2, K0R4L6 as *To*Aureo3, A0A6S9P453 as *To*Aureo4, K0TLS2 as *To*Aureo5 and A0A6S9MUW4 as *To*Aureo6. Sequence comparison between aureochromes from marine diatoms *Thalassiosira* and *Pheodactylum* reveals significant homology between *To*Aureo1/6 with *Pt*Aureo1a, *To*Aureo2 with *Pt*Aureo2, *To*Aureo3 with *Pt*Aureo1b, *To*Aureo4/5 with *Pt*Aureo1c.

### 3.2 Preliminary Characterization of Unstructured N-terminal extension of *To*AubZL

The N-terminal extension of NTE of *To*Aureo1 has hitherto remained uncharacterized. IUPred2A/AIUPRED [(Erdős & Dosztányi, 2024; Mészáros et al., 2018)] detected the presence of large stretches of Intrinsically Disordered Protein Regions (IDPRs). Several factors including binding with other protein partners or changes in pH, temperature or redox environment can influence the structure of IDPRs, disregarding the typical protein structure-function correlations. Upon binding to specific partners or changes in cellular milieu IDPRs can undergo order <-> disorder transition — often observed in gene expression or cellular signaling pathways. The ANCHOR2 prediction algorithm is used in parallel to find disordered binding regions and aids in identification of transition from unstructured to the structured states in presence of suitable protein partner. The results reveal that barring short amino acid residue stretches at the extreme N-terminal end, the whole NTE (N-Terminal Extension) of *To*Aureo1 is largely disordered. 18-53rd residue stretch is additionally redox-sensitive [**Figure S3**]. Disordered binding regions are in between 111 to 141 (precisely preceding the glutamine-rich sequence) and again between 160th residue till the end of NTE. The intrinsic disorder of IDPRs is crucial for transcriptional regulation.

EXPASY ScanProsite revealed the existence of glutamine-rich amino acid stretch from 141-171. However, this glutamine-rich stretch does not appear to participate in binding according to both IUPred2A or more advanced AIUPRED’s ANCHOR2 algorithm. Long glutamine-rich repeats are common in several important transcription factors including CREB, which are known to regulate 4000 genes from the human genome (Martinez-Yamout et al., 2023). The glutamine-rich repeats in both CREB and TFIID are known to interact with each other to facilitate transactivation. NMR experiments revealed β-hairpin structures for both the glutamine-rich repeats in CREB - placing glutamine residues in close proximity to each other. Very interestingly, AlphaFold [AF-KOSY-93] predicts a perfect helical structure for *To*Aureo1’s glutamine-rich stretch [141-171] with a mostly above average pLDDT (confidence) score. The prediction of helical structure against this stretch is further supported by another neural network training based secondary structure prediction algorithm, PSSpred. Except for the extreme N-terminal end and glutamine-rich stretches, both AlphaFold and PSSpred reveal that NTE is mostly unstructured. However, further experiments are necessary to understand whether or not and how the conformational dynamics of IDPRs in NTE mediates the transactivation process. This will be fundamental towards understanding the general mechanism of Aureos in mediating light-regulated gene expression in photosynthetic marine algae. Considering the vast and disordered nature of NTE, this region is excluded for the present *in silico* and *in vitro* studies.

### 3.2 Modeling, docking and protein-DNA interactions

After obtaining the correct sequences of *To*Aureos, *To*AubZL from *To*Aureo1 was subjected to homology modeling. Homology was not found against the linker region [**Figure S4**]; hence the LOV domain and bZIP domains were modelled separately [**Figure 1**]. LOV was modelled using Discovery studio taking 1N9L as template (LOV of *Chlamydomonas reinhardtii*). Molprobity analysis showed 97.8% of the structure falling under Ramachandran favored region. Chimera was next used for energy minimization (Pettersen et al., 2004). Energy minimized structure gave us a 100% Ramachandran favored structure with no rotamer outliers, thus providing a satisfactory model of *To*Aureo1 LOV [**Figure 1A**]. Close inspection of FMN-binding environments in different Aureos as well as *As*Phot1 LOV reveals overall sequence/structure similarity of *To*Aureo1 LOV with others [**Figure 2**]. bZIP on the other hand was modelled using SWISS-MODEL (Waterhouse et al., 2018) and PyMol following the method of Quaternary structure modeling, followed by energy minimization using Chimera. Molprobity analysis revealed a 100% Ramachandran favoured structure with no rotamer outliers. The energy minimized *To*Aureo1 bZIP, docked with ‘Aureo-box’ (TGACGT) containing CRE DNA substrate [**Figure 1B**] disclosed favorable interaction between the two. Several amino acids from the basic region [marked in bold, RRERN**R**E**H**A**KR**S**R**I**RK**KFLLE] were found to be interacting with the Aureo-box containing CRE DNA substrate [**Figure 1C**]. The corresponding nucleotides interacting with these amino acids are highlighted in orange [**Figure 1B, C**]. bZIP-DNA interactions are mostly conserved in *To*Aureo1, as seen in Aureos, especially Aureo1 from another stramenopile (Khamaru et al., 2022).

**Figure 1:**
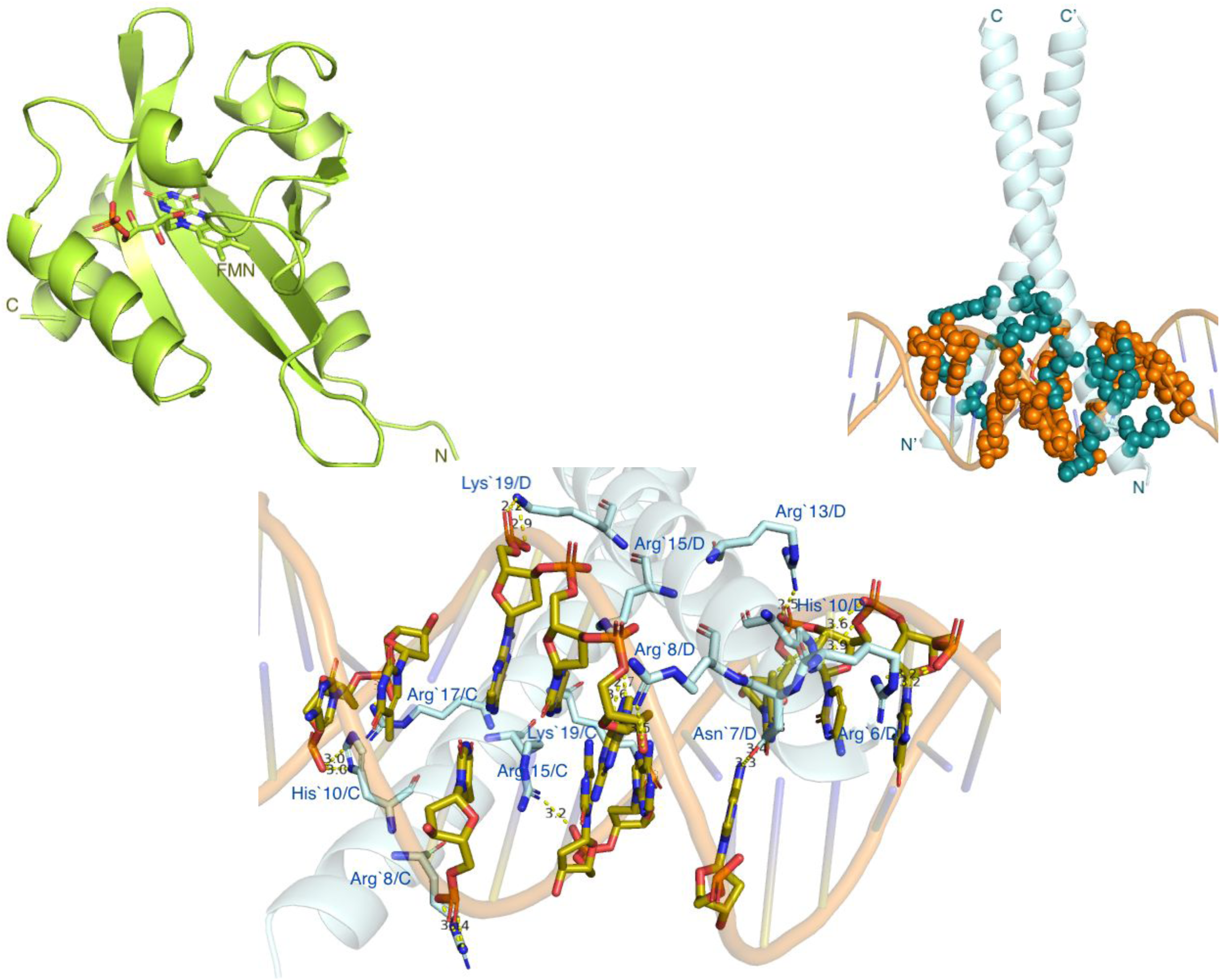
**A**: Homology modelling of *To*Aureo1 LOV domain using LOV1 domain from *Chlamydomonas reinhardtii* Phototropin (PDB ID: 1N9L) as a template shows overall conserved structure of LOV sensor. **B**: Homology modelling of *To*Aureo1 bZIP domain based on CRE binding CREB transcription factor (PDB ID: 1DH3) reveals conserved structure of bZIP. Considering few sequence alterations at the basic region, *To*Aureo1 bZIP was subjected to docking with CRE DNA substrate as obtained from PDB file 1DH3, containing the Aureo-box TGACGT. **C**: Analysis of TobZ-CRE DNA interaction shows the overall conservation of DNA binding residues at the basic region of bZIP.

**Figure 2:**
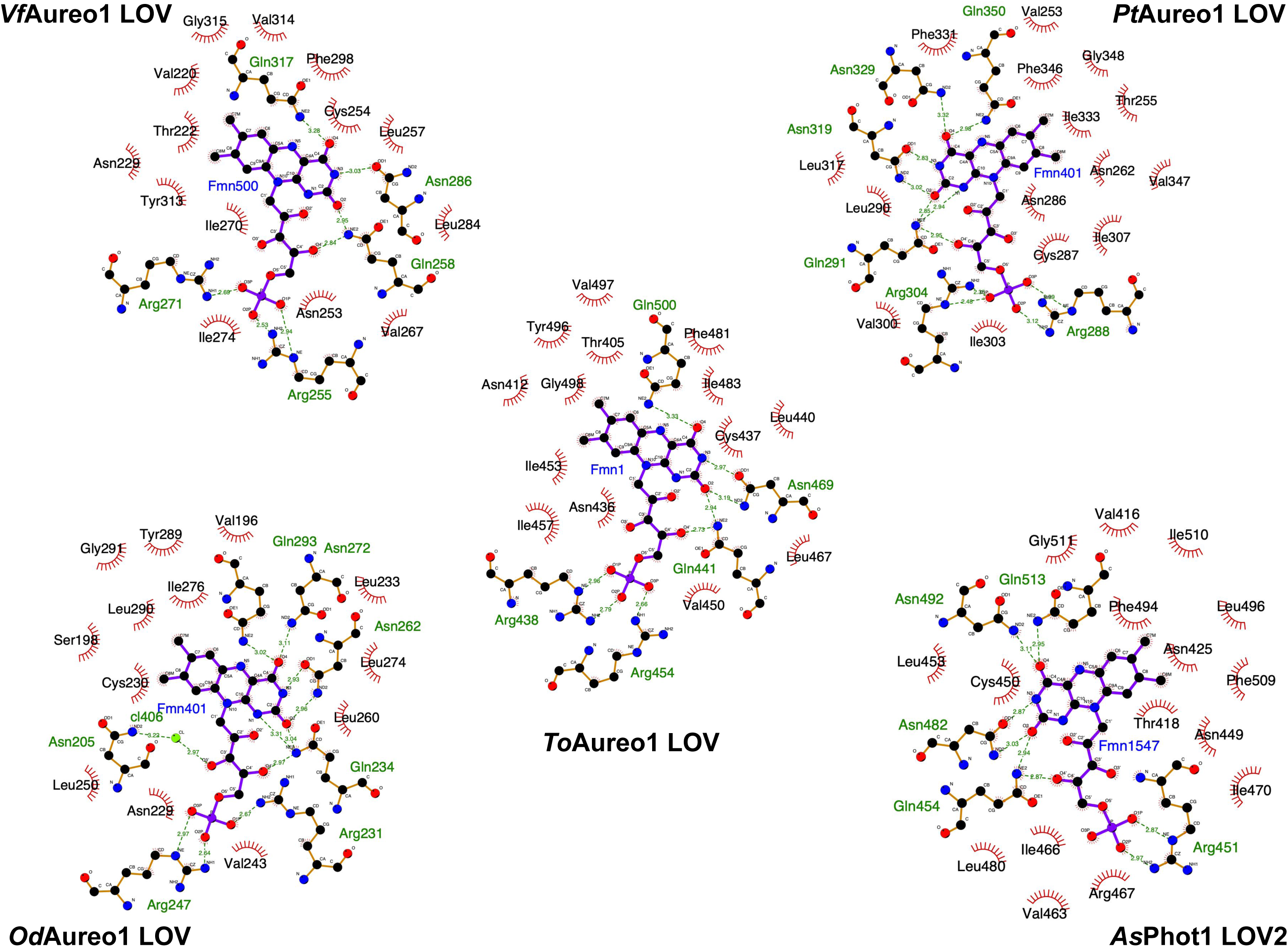
Comparison of flavin chromophore binding environment of modeled *To*Aureo1 LOV with the experimental LOV domain structures available for *Vf*Aureo1 (PDB ID: 3UE6), *Pt*Aureo1 (PDB ID: 5DKK), *Od*Aureo1 (PDB ID: 6I20) and Avena sativa phototropin1 (PDB ID: 2V0U)

### 3.3 Cloning expression and purification of *To*Aureo1 LOV and *To*AubZL

Both the constructs were successfully cloned in pET28a and verified by DNA sequencing. Following transformation in C43 (DE3) *E. coli* competent cells, the constructs were overexpressed and purified to homogeneity. Both *To*Aureo1 LOV and *To*AubZL showed similar characteristics especially in dark recovery kinetics - therefore, for final results, we pursued *To*AubZL for both kinetics and EMSA studies. In separate batches of protein purification, ToAubZL were either purified using Ni-NTA affinity column, followed by gel filtration/desalting or Ni-NTA affinity column, followed by heparin column chromatography and desalting [**Figure S5**]. The final protein concentration for all studies except CD spectroscopy, was calculated using absorbance at 447 nm and considering the molar extinction coefficient of FMN. Whether or not *To*AubZL was subjected to high salt concentration during purification did affect photocycle kinetics and EMSA results - we discuss this in the subsequent section of our manuscript.

### 3.4 Dark recovery kinetics and Circular Dichroism of *To*AubZL

*To*AubZL, purified by gel filtration chromatography was subjected to UV-Vis spectroscopy to determine kinetic parameters. *To*AubZL revealed a typical LOV spectrum indicative of a FMN-bound state with absorption maximum at 447 nm [**Figure 3A**]. OD_447nm_ values versus time plot and subsequent fitting to a first order exponential equation revealed an adduct state lifetime of 38.5 minutes and a rate constant of 4 x 10^-3^ sec^-1^.

**Figure 3:**
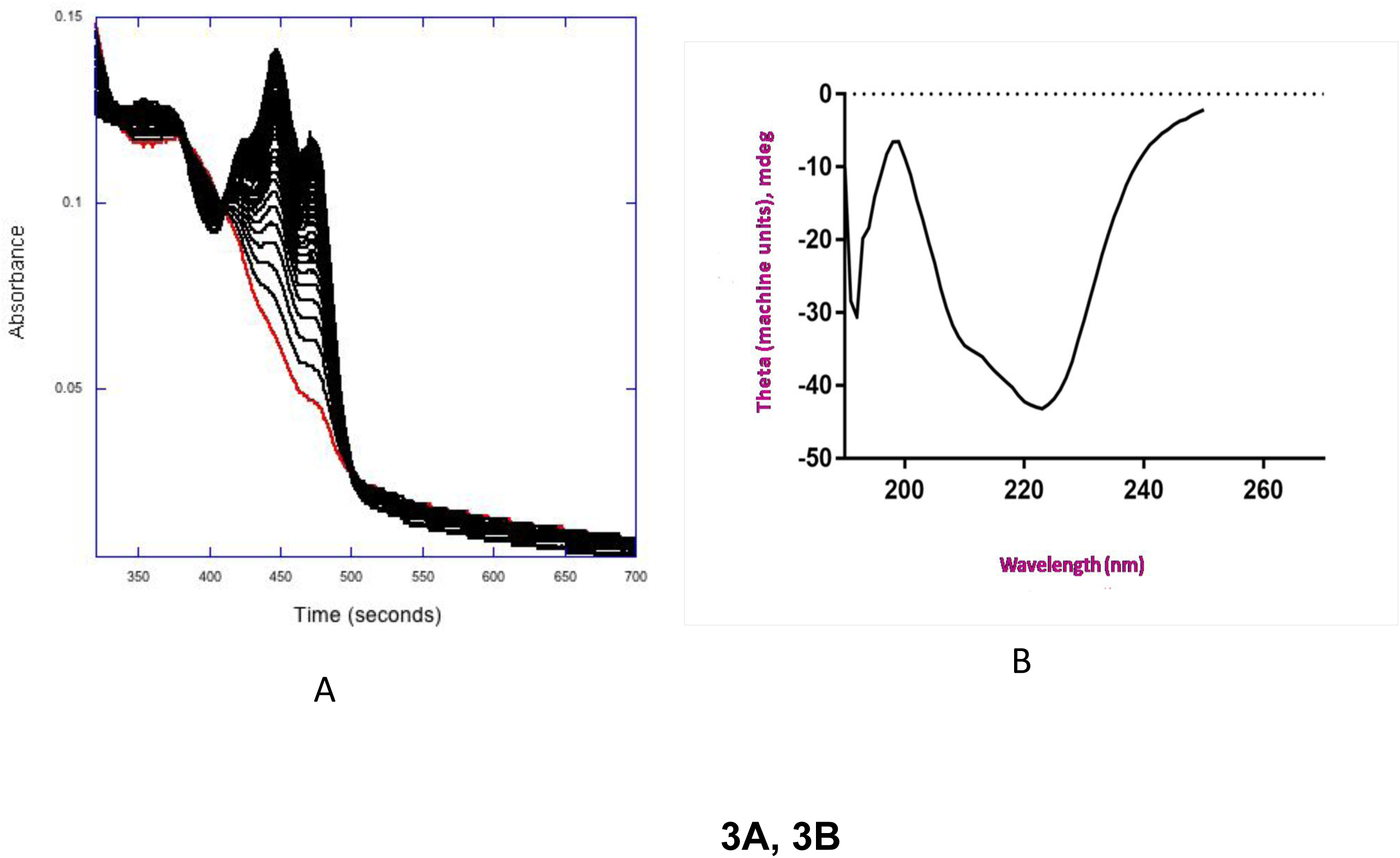
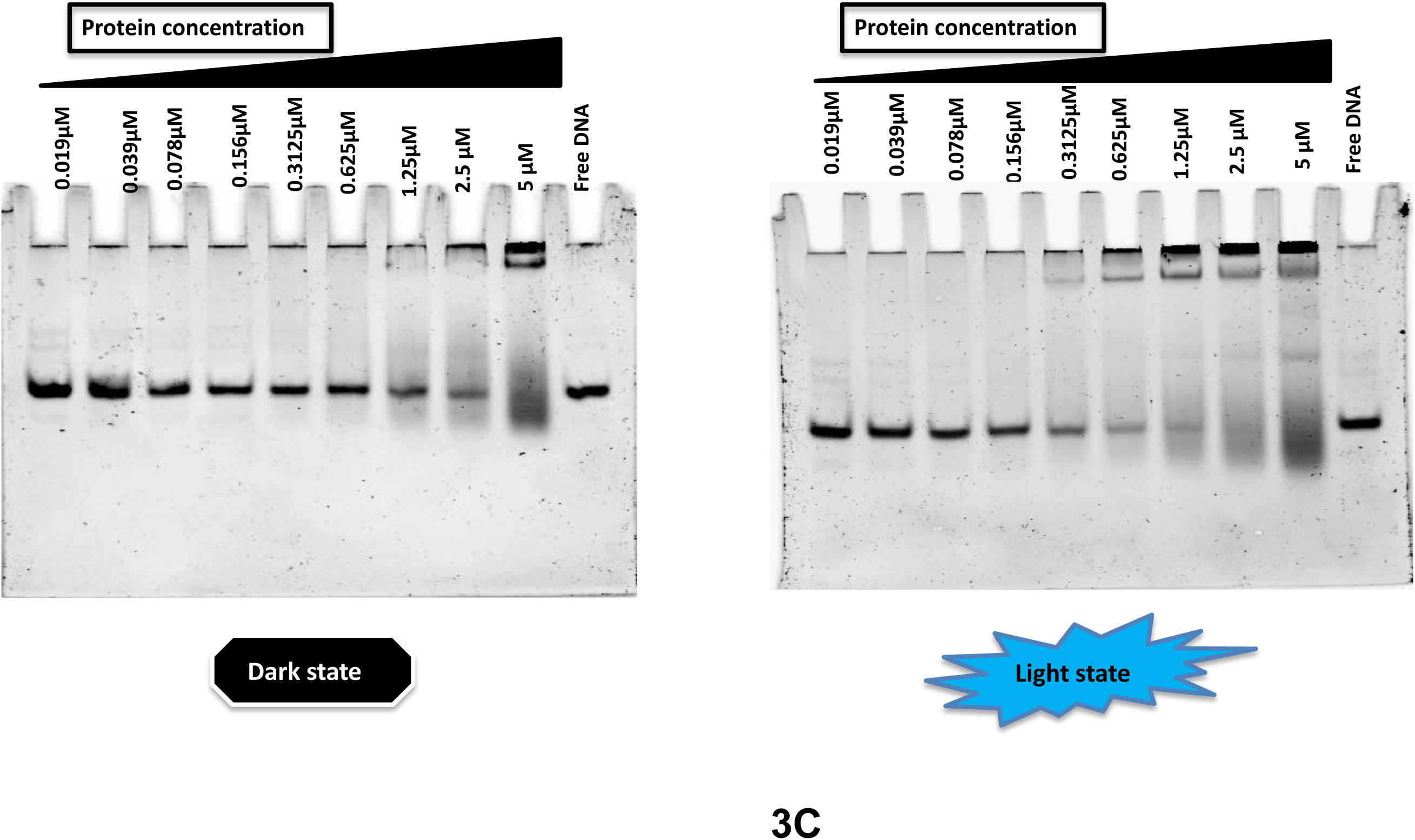
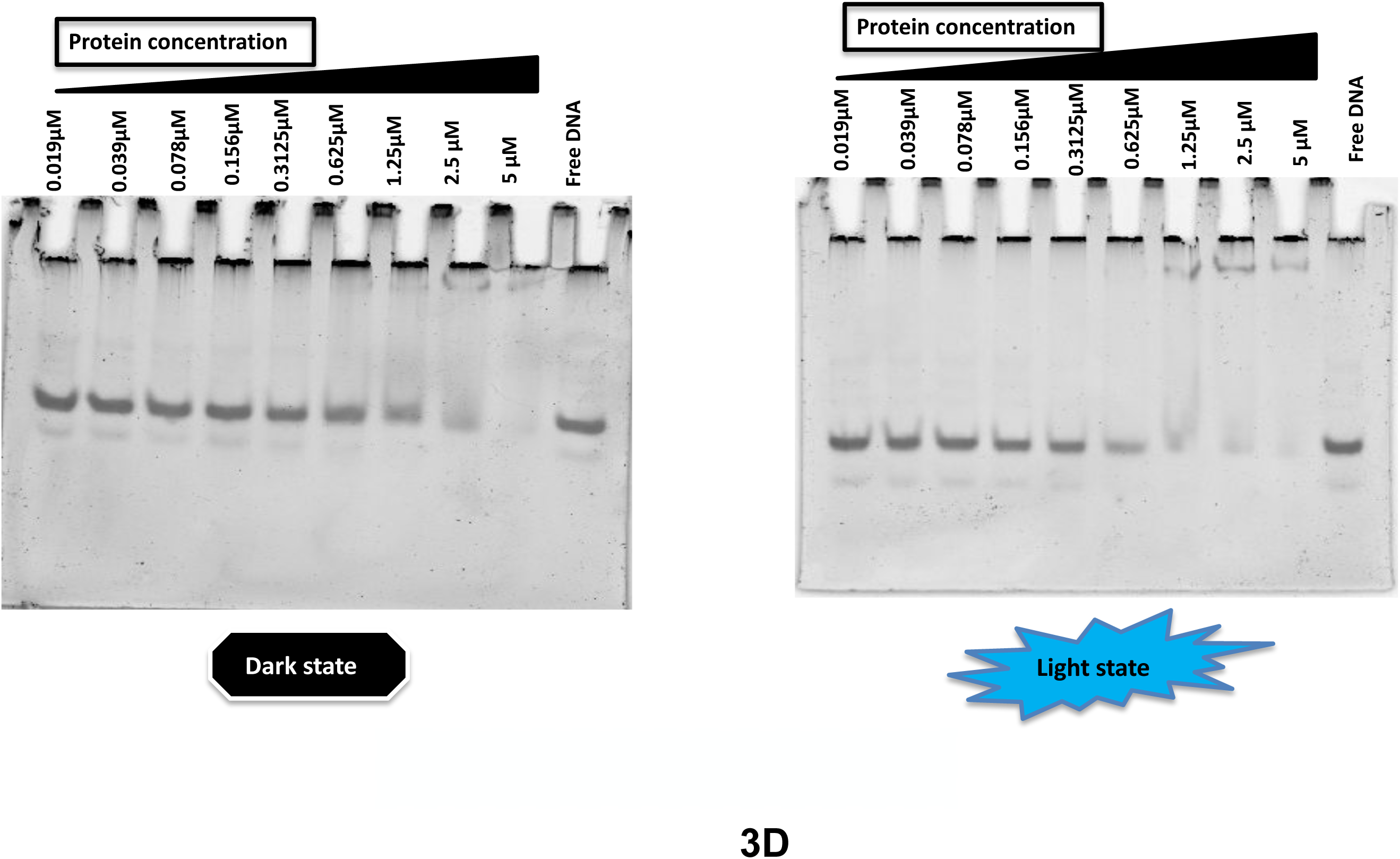
**A**: Reversible dark recovery kinetics with absorption maximum at 447nm of *To*AubZL reveals functionally active form of purified *To*AubZL. However, *To*AubZL shows a relatively slow photo cycle with a half life of 38.5 minutes and rate constant 4 x 10^-3^ s^-1^. **B**: CD spectroscopy reveals high content of beta and flexible loop structures in *To*Aureo1 with low percentage of alpha-helical structures. **C and D**: Electrophoretic mobility shift assay (EMSA), performed with the Aureo-box containing substrate DNA and *To*AubZL. EMSA studies were conducted from multiple batches of *To*AubZL purification [Purified maintaining low salt conditions - C; purified after high salt exposure during Heparin chromatography - D]. *To*AubL, very clearly showed 4 fold higher affinity binding under *in vitro* light conditions, especially when purified through low salt conditions.

Far-UV (190-250 nm) CD spectroscopy was conducted with purified *To*AubZL at a concentration of 1.9 mg/ml (concentration was calculated using absorbance at 280 nm). *To*AubZL CD spectrum [**Figure 3B**] revealed high percentage of beta sheet structures – anti-parallel beta sheets with beta bulges is the core of LOV domain. The presence of the bZIP domain at the N-terminus and the short helical linkers at the LOV domain is reflected with alpha helices, albeit in low amounts. Except for alpha helices, the predictions of secondary structural elements from both the algorithms as listed in the table corroborate each other well [**Table S1-A**]. The high percentage of disordered regions in both the WT and synthetic constructs indicate the long, flexible linker region connecting the sensor and effector domains. Besides the flexible linker, the bZIP domain, especially at the basic DNA binding region, might also exist in a disordered state prior to DNA binding (Podust et al., 2001). In fact, fly-casting theory (Shoemaker et al., 2000) suggests that disorderliness of the basic region accelerates DNA binding kinetics. A recent study, however, reveals that the dimerization domain (zipper region) remains in a largely ordered conformation in both unbound as well as transition states. However, dimerization introduces additional helical propensity at the basic region, which results in a fully folded conformation upon DNA binding (Kuravsky et al., 2024).

### 3.5 Electrophoretic mobility shift assay

*To*AubZL purified maintaining low salt conditions (Ni-NTA + gel filtration/desalting maintaining 50 mM NaCl concentration) showed prominent increase in DNA binding activity under blue-light [**Figure 3C**]. Super-shifted DNA is clearly visible from 312.5 nM, whereas the same appears from 1.25 µM onwards in the dark state. However, when purified by Ni-NTA + desalting + Heparin + desalting and thus exposed under high salt concentration [**Figure 3D**] during heparin chromatography, the difference in DNA binding affinity decreases. In this case, we observe marginal increase in DNA binding affinity under light state. This result is consistent with our finding that *To*AubZL, when purified through high salt exposure, departs from typical dark recovery kinetics after a certain time. After the protein starts recovering and maintains the normal course of dark recovery for a shorter period of time, the overall absorbance across the entire scan-range starts increasing significantly -- indicating possible change of conformation/unfolding at the flavin binding site. *To*AubZL shows similar characteristics of light-dependent DNA binding like it’s counterpart *Pt*Aureo1 [(A. Banerjee, Herman, Serif, et al., 2016; Schellenberger Costa et al., 2013)]. It is perhaps worth-mentioning that both *Phaeodactylum* and *Thalassiosira* Aureo1 show light dependent DNA binding in presence of MgCl_2_.

### 3.6 Importance of light-regulated network motifs for transcriptional regulation - Design and engineering of *AsTo*bZL

Figure 4 depicts two major types of biological circuit design strategies prototyping “AND” and “OR” logic gates. In “AND” operation, we get an output ‘Z’ only when both ‘A’+ ‘B’ are activated, while in “OR” gate activation of ‘A/B’ leads to output ‘Z’. Therefore the initiation time of the “AND” gate is higher than that of “OR” gate. For the same reason, in the reverse scenario (upon removal of the light source) “AND” gate is switched off before the “OR” gate. For generating sensory transcriptional regulation networks (STRN), it is possible to add additional layers of control via introducing environmental stimuli in the input function. Non-toxicity of visible light (especially in low doses), high quantum efficiency (S. Banerjee & Mitra, 2020) as well as varied light-state lifetime of photo-reversible LOV sensors, make LOV-based TFs an ideal choice for developing STRN/light regulated gene expression circuits. To elaborate the importance of light/adduct-state lifetime on STRN, we consider a hypothetical scenario [Figure 4]. In this example, the activation threshold of photoreceptor “A” is ‘t’ seconds while that of B is ‘s’ seconds. Now, in a double-input circuit, the formation of product ‘Z’ will be delayed in “AND” and not delayed in “OR” operations by ‘s’ seconds. The time for photoactivation being similar for most LOV sensors, the activation part is less complicated. During dark-reversal, let’s consider that the deactivation thresholds of “A” and “B” are ‘e’ and ‘l’ seconds respectively. In this case, the “AND” circuit will be switched off faster, immediately after dark reversal of “A”. However, the “OR” circuit will be delayed further by ‘l’ seconds. Now, the time ‘e’, known as dark recovery time or light/adduct-state lifetime, varies from one photoreceptor to another. Moreover, the motifs of STRN are temporally precise. Therefore, by choosing a suitable LOV variant, it is possible to fine tune the output of these circuit operations and achieve the results with temporal precision.

**Figure 4:**
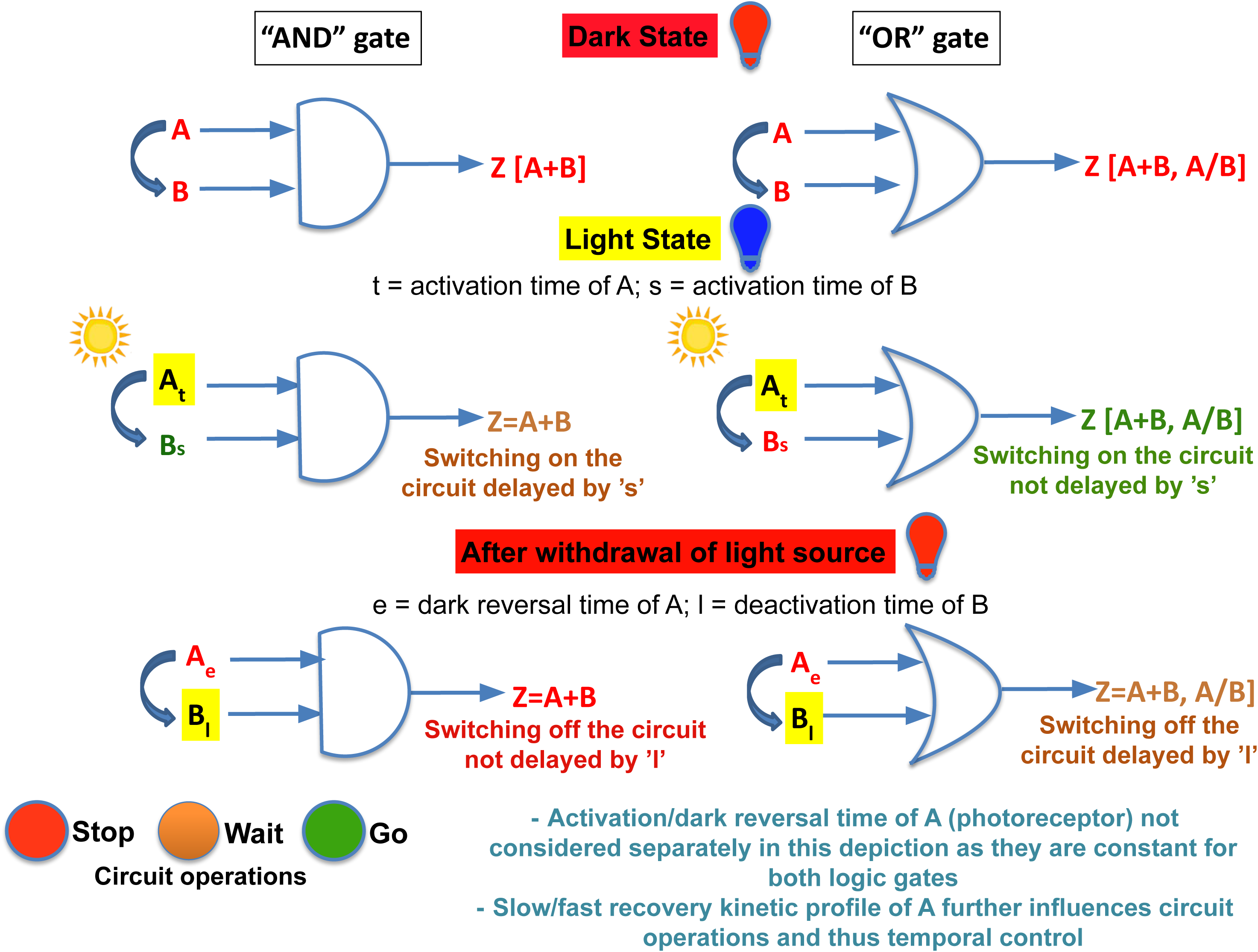
Schematic representation of light-mediated synthetic logic circuit and importance of dark reversal time of photoreceptor in manipulating the outputs from synbio circuit. Aureochromes being photoreceptor-cum-transcription factors can carry out temporal expression of downstream genes differentially under dark and light conditions. The pattern of gene expression would essentially depend on a) dark reversal time of the LOV sensor, b) DNA binding affinity of the effector bZIP and c) nature of the logic gate under operation.

On this account and also to understand the importance of heterogeneous linker [**Figure S4**] connecting sensor and effector, we designed and constructed *AsTo*bZL. Another reason behind the selection of *As*LOV2 as sensor is that this is the most well studied and preferred LOV sensor for designing optogenetic constructs (S. Banerjee & Mitra, 2020). It is worth mentioning here that the linker region in between effector and sensor is poorly conserved in different LOV photoreceptors. This region varies in sequence and length even within the same family, e.g. Aureos [**Figure S4**]. However, it is obvious that this flexible and variable linker is essential for transducing signal from sensor to the effector. But then the question is whether the much shorter linker of *Avena sativa (As)* Phot1 with completely different sequence composition is suitable for designing light-regulated TF. We therefore constructed *AsTo*bZL combining the N-terminal bZIP domain from *To*Aureo1 and linker + LOV2 components from *As*Phot1 [Figure 5A].

**Figure 5:**
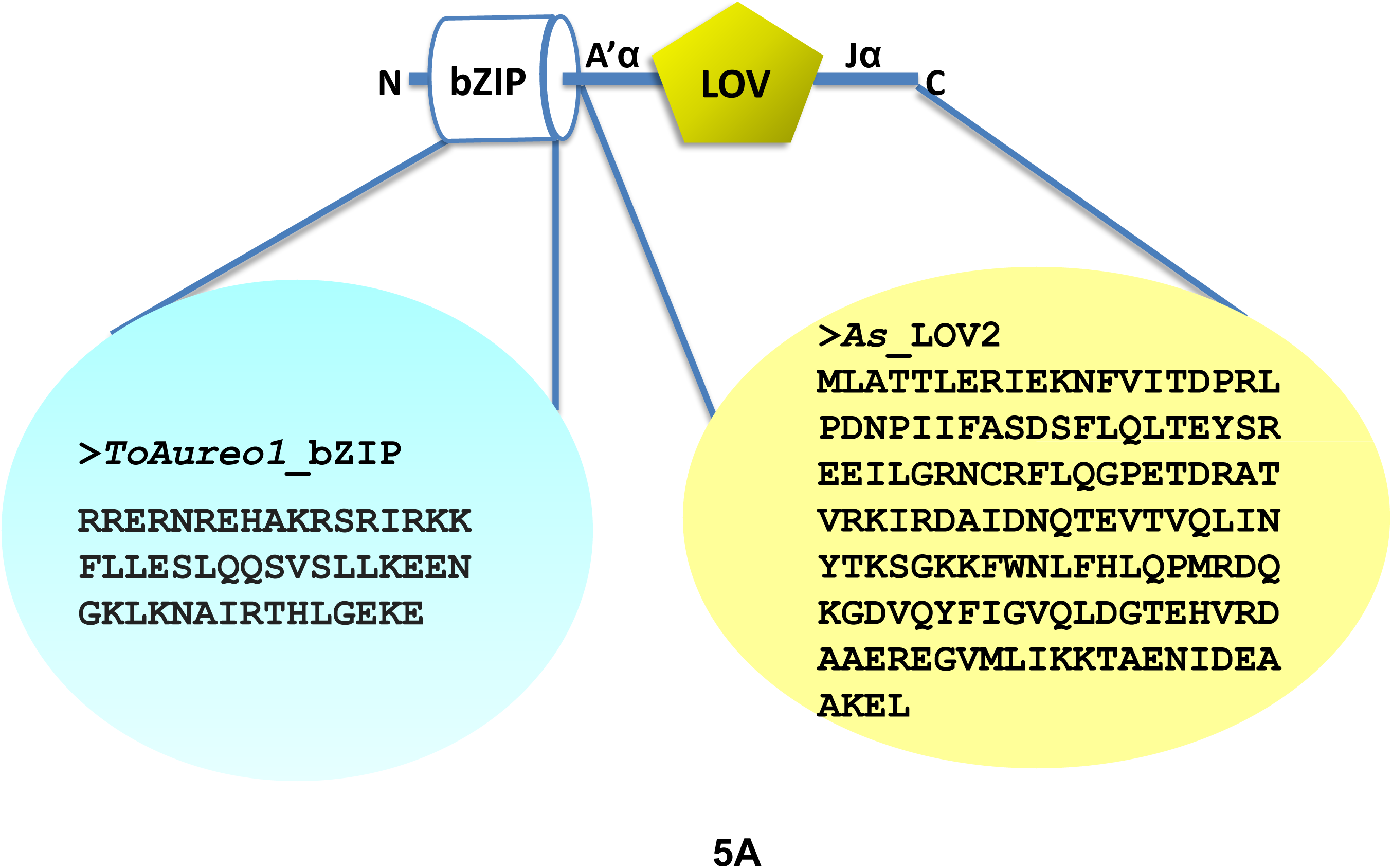
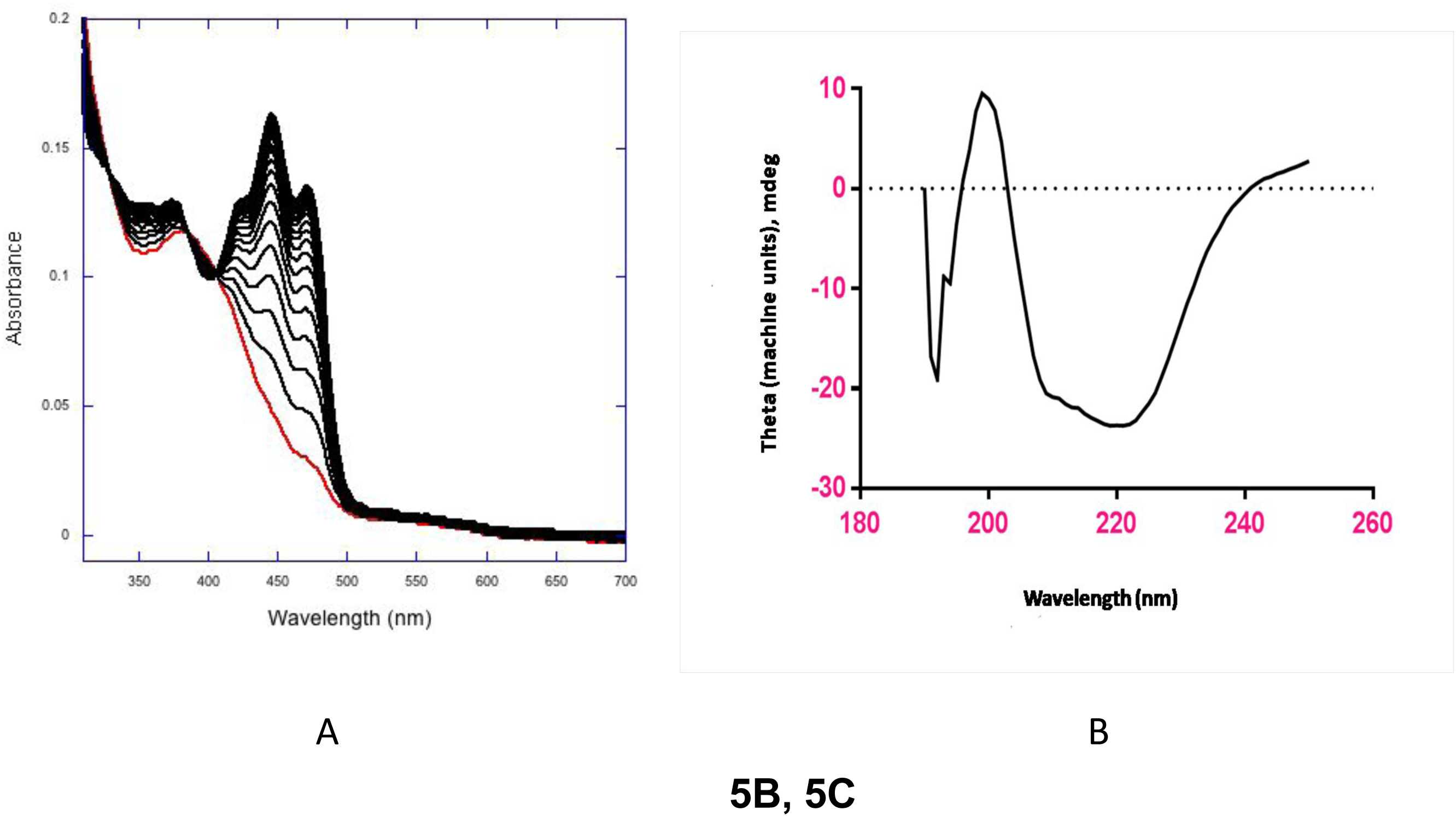
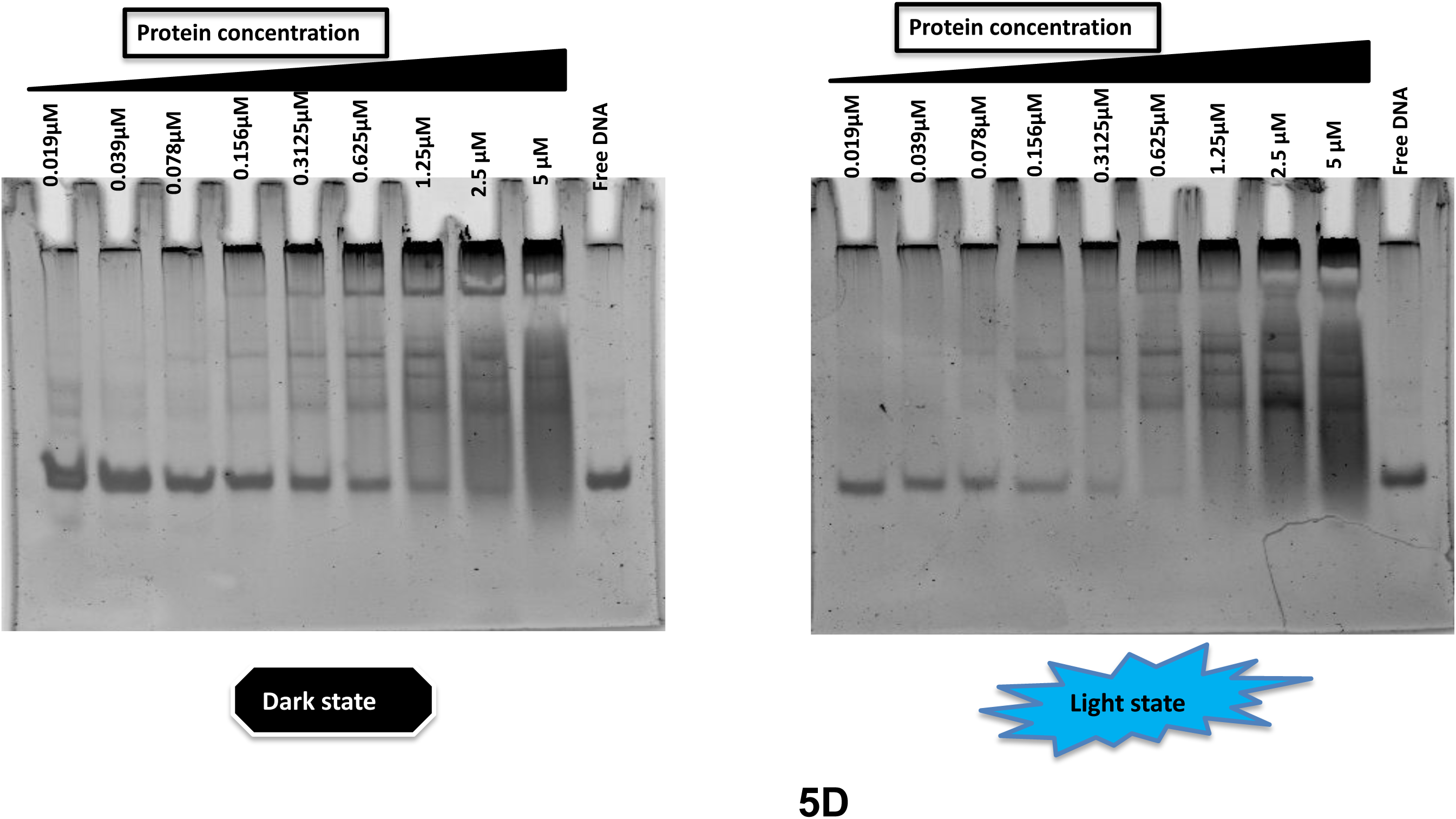
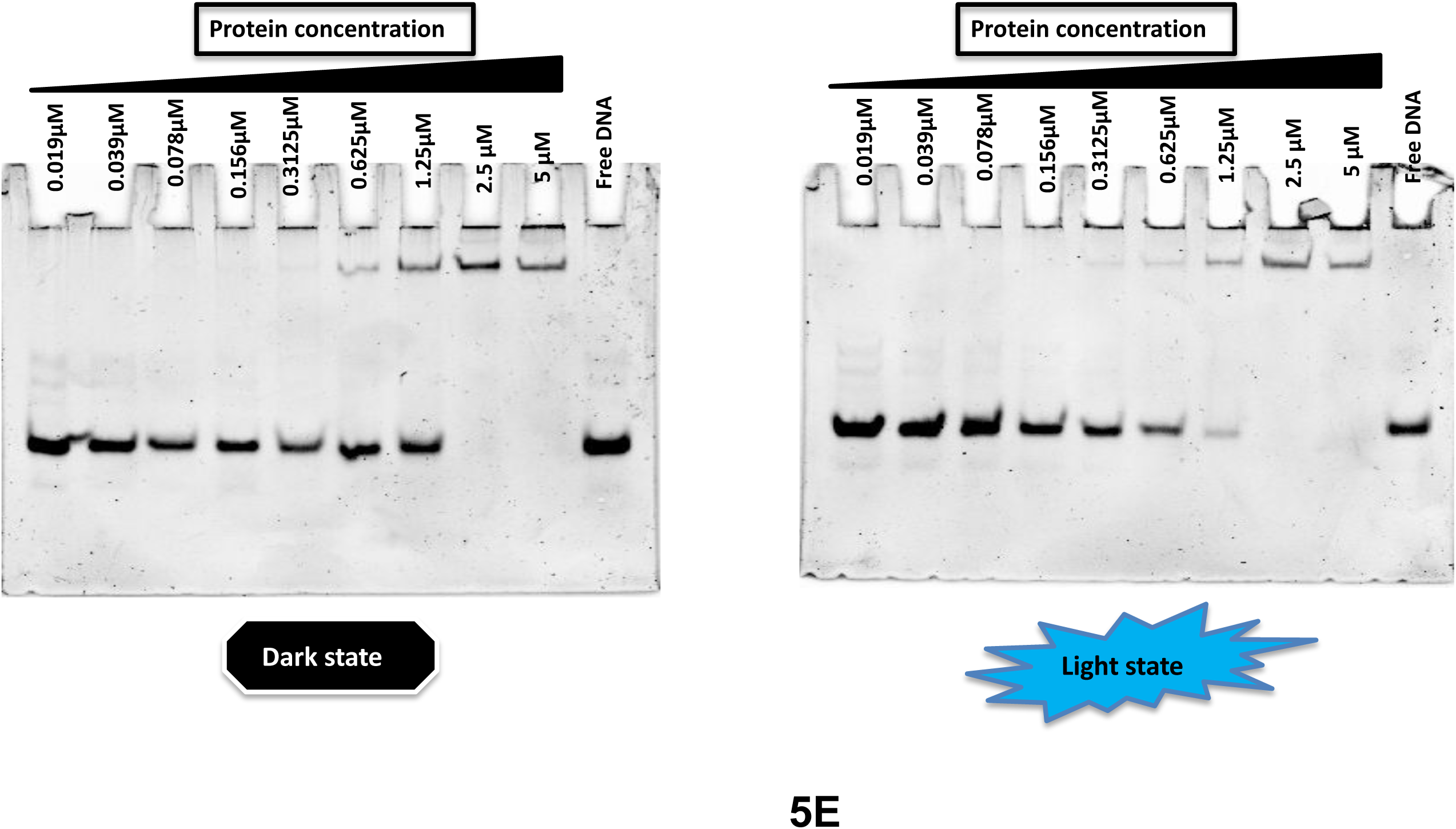
**A**: Construction of synthetic bZIP-LOV construct, using N-terminal bZIP sequence from *To*Aureo1, shorter linker + LOV domain sequences from heavily used optogenetic photoreceptor, Phototropin LOV (*As*LOV2) from *Avena sativa*. This construct is created to check the a) utility of shorter vs longer linker sequences in *As*Phot1 vs *To*Aureo1; and b) impact of exchanging LOV sensor on the DNA binding activity under light-dark conditions. **B**: Reversible dark recovery kinetics with absorption maximum at 447 nm of *AsTo*bZL reveals functionally active form of purified *AsTo*bZL. As expected, *AsTo*bZL shows faster photo cycle kinetics with a half life of 231 seconds and rate constant 1 x 10^-2^ s^-1^. **C**: CD spectroscopy reveals similarity in secondary structure content between *AsTo*bZL with *To*AubZL. **D and E**: Electrophoretic mobility shift assay (EMSA) was performed with the Aureo-box containing substrate DNA. EMSA studies were conducted from multiple batches of *AsTo*bZL purification [Purified maintaining low salt conditions - D; purified after high salt exposure during Heparin chromatography - E]. Similar to *To*AubZL, *AsTo*bZL too showed 4 fold higher affinity binding under *in vitro* light conditions, especially when purified through low salt conditions.

### 3.7 Cloning, Expression and Characterisation of *AsTo*bZL

Following fusion PCR, *AsTo*bZL was successfully cloned into pET28a vector and transformed to C43 (DE3) strain of *E. coli* for protein production. *AsTo*bZL was purified to homogeneity [**Figure S5**] using the same purification protocol as *To*AubZL *i.e.* maintaining low and high salt conditions. Incidentally, *AsTo*bZ also showed similar trends in both UV-Vis and EMSA studies. The reversible dark recovery kinetics of *AsTo*bZL [Figure 5B] completed on a much faster time scale (lifetime 231 seconds) as expected, with a rate constant of 1 x 10^-2^ sec^-1^ – comparable to the photocycle kinetics of wild type *As*LOV2 (Zoltowski et al., 2009). CD spectroscopy, conducted at 0.88 mg/ml concentration, revealed a high percentage of beta strands and disordered regions with small fraction of alpha helices, similar to *To*AubZL [Figure 5C**, Table S1-B**]. Overall, the CD statistics of both *To*AubZL and *AsTo*bZL are quite similar. In EMSA studies, when purified under low salt conditions, free DNA disappears from 625 nM onwards, while the same could be visible till 2.5 µM in the dark state. *AsTo*bZ purified through heparin chromatography showed less than two fold differences in DNA binding affinity between dark and light conditions. Success of *AsTo*bZL as blue-light responsive TF indicates that in presence of a suitable linker (in this case, A’α of *As*LOV2), even with completely different amino acid residue composition and shorter length, it is possible to design efficient optogenetic constructs. Additionally, replacing modular LOV sensor with varying light-state lifetime can allow us to fine tune the target physiological phenomenon.

### 3.8 Investigation on probable higher order structures in Aureos and implications in signaling

Aureochromes being blue-light responsive photoreceptor cum TFs are intricately involved in the light signaling mechanism of photosynthetic stramenopiles. As mentioned earlier in our manuscript, *Phaeodactylum tricornutum* Aureos (*Pt*Aureos) are perhaps the most well studied Aureos not only among the diatoms but in all stramenopiles. It is remarkable that a single photoreceptor-TF like *Pt*Aureo1a can influence the expression of 75% genes upon transition from red- to blue-light (Mann et al., 2020). And, disruption of *Pt*Aureo1a could almost eliminate light-induced signaling in this diatom. Not just this, another paralog *Pt*Aureo1c enables *Phaeodactylum* to get acclimatized under very high light conditions by promoting non-photochemical quenching required for photo-protection (Zhang et al., 2024). Such unique ability of photo-adaptation places diatoms as the leading primary producers in the coastal region, ahead of green algae. Therefore, it is clear that *Pt*Aureo paralogs can influence global gene expression – the question is how? As per recent *in vivo* and *in vitro* experimental evidence (Im et al., 2024), *Pt*Aureos can carry out differential gene expression through homo/heterodimerization in all possible combinations. The formation of dimer/higher order structures in *Pt*Aureos has been established under blue-light conditions using size exclusion chromatography (Zhang et al., 2024). Recently, we have also depicted the homo/heterodimerization potential of Aureochromes from *Ectocarpus sp*. (Khamaru et al., 2024) using structural bioinformatics and information-theoretic approach. Herein, we therefore wanted to check whether *To*Aureos too has the homo/heterodimerization potential. The helical wheel projection [Figure 6] that explores the heterodimerization potential of *To*Aureo1 (the major focus of this manuscript) reveals that the ‘a’ and ‘d’ positions of the heptad repeats in all Aureos are predominantly occupied by non-polar amino acids, especially leucine. Furthermore, the ‘e-g′’, or ‘e′-g’ positions make favourable electrostatic interactions among all the *To*Aureos - indicating stable bZIP dimers and compatibility among all *To*Aureo hetero-as well as homodimers [**Figure S6**].

**Figure 6:**
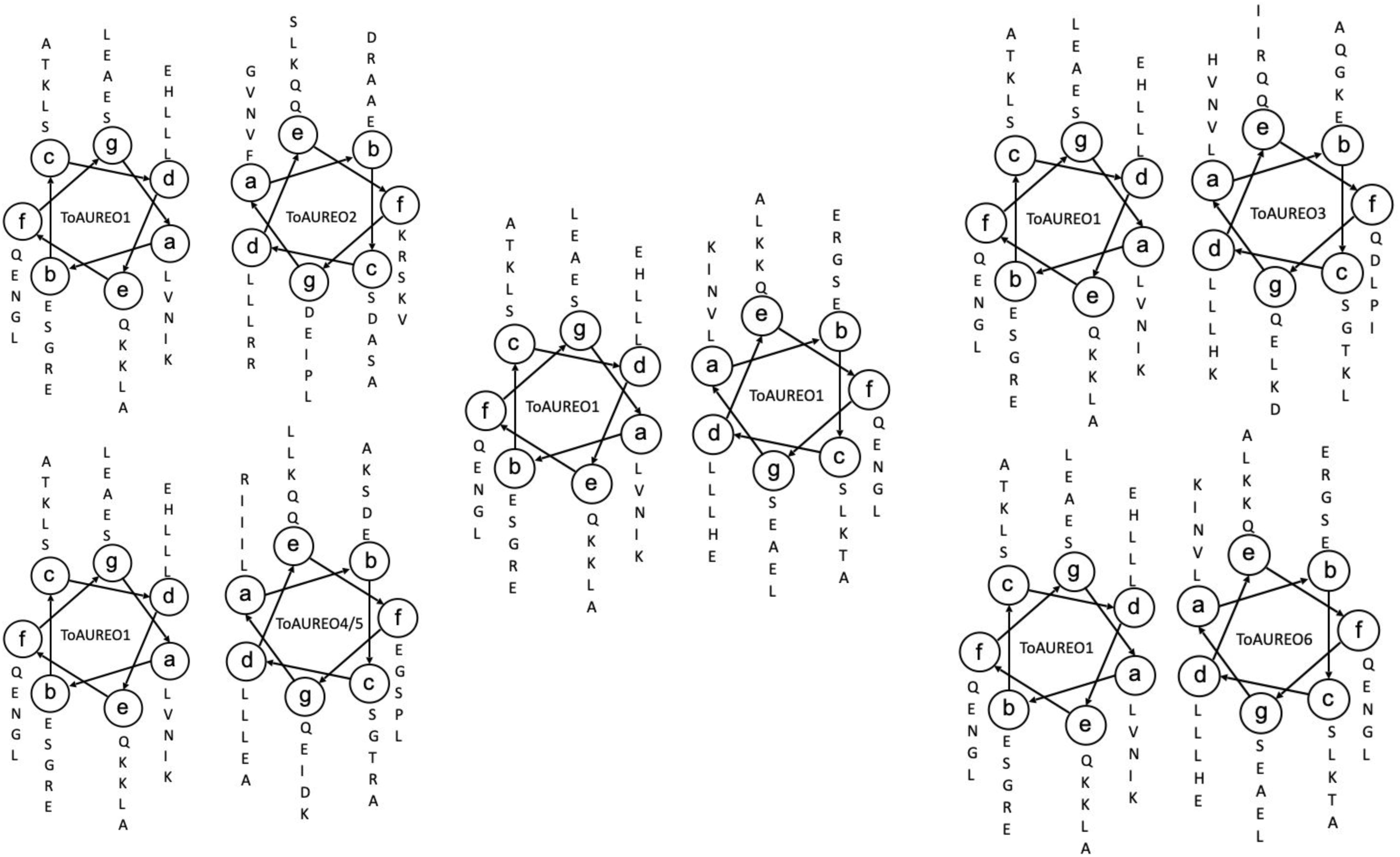
Helical wheel projections showing the compatibility of *To*Aureo1 with other *To*Aureo sequences towards stable hetero-dimer interactions with consequences in DNA binding activities and gene expression.

## 4. Discussion

Aureos are exclusively present in photosynthetic heterokonts. Consisting of yellow-green algae, diatoms, brown algae etc., these photosynthetic algae inhabit different strata of marine environment and exhibit diverse physiological responses upon blue-light irradiation. Interestingly, Phototropins, which are the major blue-light sensitive LOV photoreceptors in land plants, are not found in photosynthetic heterokonts where Aureochromes are present. The diatom *Thalassiosira oceanica* was once considered identical to *Thalassiosira pseudonana,* though later it was found to possess a much larger genome size of ~80MB. Significantly large size of *T. oceanica* from both of its closely related species, *Thalassiosira pseudonana* (~34 MB) and *Phaeodactylum tricornutum (Pt)* (~28 MB) is attributed to its DNA recruitment from multiple sources (Lommer et al., 2012). Although the exact function of *To*Aureo1 is not known, *Pt*Aureo1(a/b/c) and *Pt*Aureo2 have been studied in great detail with extensive crystallography studies at different dark/light conditions [(A. Banerjee, Herman, Kottke, et al., 2016b; Heintz & Schlichting, 2016)] and physiological responses manifested by light-regulated transcriptional network [(Coesel, 2024; Im et al., 2024)]. *Pt*Aureo1a is also found to be important in regulating the expression of cell cycle protein dsCyc2, which facilitates the transition of a cell from G1 checkpoint (Huysman et al., 2013). LOV domain of *Pt*Aureo1 upon blue-light absorption unfolds its flanking alpha helices in a similar pattern as phototropin LOV which in turn induces LOV dimerization (Herman et al., 2013). This phenomena can be critical in blue-light dependent DNA binding (Kobayashi et al., 2020). *To*AubZL or its synthetic counterpart *AsTo*bZL can behave in a similar way. As discussed in the results section, the destabilization of LOV sensors following high salt exposure did impact DNA binding by *To*AubZL/*AsTo*bZL. Similarly in *Vf*Aureo1 bZIP+LOV (photozipper), conformational changes in flavin binding residues in LOV domain influences blue-light mediated DNA binding by bZIP (Kobayashi et al., 2020). We explain this further by TF cooperativity.

TF cooperativity involves multitude of events and is imperative to attain DNA-binding specificity across myriad TFs families (Badis et al., 2009). TFs generally bind to their cognate DNA substrates with the aid of relatively shorter DNA binding domains. However, TFs often possess long/short stretches of IDPRs either at the N or C-terminal to the DNA binding domain. IDPRs are essential for transcriptional regulation that includes binding with the chromatin, recruitment of cofactors and coactivators - thus assembling the transcriptional machinery. While IDPRs do have an impact on binding to particular target sites and thereby facilitating DNA binding interactions [(Brodsky et al., 2020, 2021; Jolma et al., 2013)], it is still not well characterized what causes this DNA binding specificity. TFs temporarily bind to DNA non-specifically in order to search the genome for specific DNA sequences so that they can bind with greater affinity. In the facilitated diffusion model, the TF diffuses arbitrarily until it finds a nonspecific binding site, which generally involves short-lived binding events. In hopping and sliding methods, exchanging between one and three dimensional diffusion processes, TFs firmly and specifically bind to the target site on the genome – considered as long-lasting binding events. In essence, TFs weakly binds to the DNA to mediate faster protein diffusion in search mode and upon finding target site i.e. in recognition mode it permits the stronger and stable TF-DNA interactions (Al Masri et al., 2023).

Now the questions are: what is the precise role of light in TF binding and gene expression? Can light promote/facilitate non-specific to specific binding of TF to DNA? Non-specific interactions are predominantly electrostatic in nature, whereas binding with the substrate involves more base specific interactions like hydrogen bonding and van der Waals’ interactions [(Al Masri et al., 2023; De Jonge et al., 2022; Sang & Johnson, 2024)]. Nevertheless, switching from non-specific to specific binding involves conformational changes of TF and/or DNA. It is a well known fact that light triggers conformational changes in the LOV sensor and a signal is generated. The signal is then transduced to the DNA binding domain of the TF to bind with DNA. It is further possible that dimerization between LOV sensors in turn promotes self-clustering of TFs resulting in enhanced DNA binding. At the genome level, TFs carry out extensive search processes mediated by non-specific interactions followed by specific interactions with the substrate DNA. Apparently, under *in vitro* conditions, light triggers binding with the substrate DNA only when bound with non-specific ones i.e. host cellular DNA. Switching between non-specific to specific binding mode greatly depends on the similarity of the amino acid residue stretches involved in the non-specific/specific protein-DNA interactions. Higher the overlap, faster is the switching and vice versa. Light probably facilitates this switching event and improves the response time. At the *in vivo* level, this can be perceived as light driven acceleration of the search process to find specific binding sites. However, this remains to be validated with further experiments.

## Acknowledgements

AD acknowledges the Department of Biotechnology (DBT), Govt of India, for her doctoral fellowship. Preliminary results of this manuscript were provided in her doctoral thesis. PS acknowledges the Council of Scientific and Industrial Research (CSIR), Govt of India, for her doctoral fellowship. DM acknowledges funding from DBT (BT/PR26435/BRB/10/1627/2017) during the initial phase and Anusandhan National Research Foundation [ANRF grant no CRG/2023/006463, formerly SERB], Govt of India during the current phase of this research. The authors also gratefully acknowledge central instrumentation facilities, developed from DBT-BUILDER and DST-FIST funds. Special thanks to fellow lab member Aparna Boral for her technical assistance during initial EMSA studies.

## Authors’ contribution

***Anwesha Deb*** - Methodology, validation, formal analysis, investigation, data curation, writing (original draft) and visualization; ***Piu Sarkar*** - Methodology, validation, formal analysis, investigation and visualization; ***Devrani Mitra*** - Conceptualization, Methodology, validation, formal analysis, investigation, data curation, writing (original draft), writing (review and editing), visualization, supervision, project administration and fund acquisition.

## References

Al Masri, C., Wan, B., & Yu, J. (2023). Nonspecific vs. Specific DNA binding free energetics of a transcription factor domain protein. Biophysical Journal, 122(22), 4476–4487. 10.1016/j.bpj.2023.10.025

Badis, G., Berger, M. F., Philippakis, A. A., Talukder, S., Gehrke, A. R., Jaeger, S. A., Chan, E. T., Metzler, G., Vedenko, A., Chen, X., Kuznetsov, H., Wang, C.-F., Coburn, D., Newburger, D. E., Morris, Q., Hughes, T. R., & Bulyk, M. L. (2009). Diversity and Complexity in DNA Recognition by Transcription Factors. Science, 324(5935), 1720–1723. 10.1126/science.1162327

Banerjee, A., Herman, E., Kottke, T., & Essen, L.-O. (2016a). Structure of a Native-like Aureochrome 1a LOV Domain Dimer from Phaeodactylum tricornutum. Structure, 24(1), 171–178. 10.1016/j.str.2015.10.022

Banerjee, A., Herman, E., Kottke, T., & Essen, L.-O. (2016b). Structure of a Native-like Aureochrome 1a LOV Domain Dimer from Phaeodactylum tricornutum. Structure, 24(1), 171–178. 10.1016/j.str.2015.10.022

Banerjee, A., Herman, E., Serif, M., Maestre-Reyna, M., Hepp, S., Pokorny, R., Kroth, P. G., Essen, L.-O., & Kottke, T. (2016). Allosteric communication between DNA-binding and light-responsive domains of diatom class I aureochromes. Nucleic Acids Research, 44(12), 5957–5970. 10.1093/nar/gkw420

Banerjee, S., & Mitra, D. (2020). Structural Basis of Design and Engineering for Advanced Plant Optogenetics. Trends in Plant Science, 25(1), 35–65. 10.1016/j.tplants.2019.10.002

Boyden, E. S., Zhang, F., Bamberg, E., Nagel, G., & Deisseroth, K. (2005). Millisecond-timescale, genetically targeted optical control of neural activity. Nature Neuroscience, 8(9), 1263–1268. 10.1038/nn1525

Brodsky, S., Jana, T., & Barkai, N. (2021). Order through disorder: The role of intrinsically disordered regions in transcription factor binding specificity. Current Opinion in Structural Biology, 71, 110–115. 10.1016/j.sbi.2021.06.011

Brodsky, S., Jana, T., Mittelman, K., Chapal, M., Kumar, D. K., Carmi, M., & Barkai, N. (2020). Intrinsically Disordered Regions Direct Transcription Factor In Vivo Binding Specificity. Molecular Cell, 79(3), 459–471.e4. 10.1016/j.molcel.2020.05.032

Caamaño, A. M., Vázquez, M. E., Martínez-Costas, J., Castedo, L., & Mascareñas, J. L. (2000). A Light-Modulated Sequence-Specific DNA-Binding Peptide. Angewandte Chemie, 39(17), 3104–3107. 10.1002/1521-3773(20000901)39:17<3104::AID-ANIE3104>3.0.CO;2-0

Chaves, I., Pokorny, R., Byrdin, M., Hoang, N., Ritz, T., Brettel, K., Essen, L.-O., Van Der Horst, G. T. J., Batschauer, A., & Ahmad, M. (2011). The Cryptochromes: Blue Light Photoreceptors in Plants and Animals. Annual Review of Plant Biology, 62(1), 335–364. 10.1146/annurev-arplant-042110-103759

Christie, J. M., Salomon, M., Nozue, K., Wada, M., & Briggs, W. R. (1999). LOV (light, oxygen, or voltage) domains of the blue-light photoreceptor phototropin (nph1): Binding sites for the chromophore flavin mononucleotide. Proceedings of the National Academy of Sciences, 96(15), 8779–8783. 10.1073/pnas.96.15.8779

Coesel, S. N. (2024). More than a photoreceptor: Aureochromes are intrinsic to the diatom light-regulated transcriptional network. Journal of Experimental Botany, 75(7), 1786–1790. 10.1093/jxb/erae004

Crosson, S., & Moffat, K. (2001). Structure of a flavin-binding plant photoreceptor domain: Insights into light-mediated signal transduction. Proceedings of the National Academy of Sciences, 98(6), 2995–3000. 10.1073/pnas.051520298

De Jonge, W. J., Patel, H. P., Meeussen, J. V. W., & Lenstra, T. L. (2022). Following the tracks: How transcription factor binding dynamics control transcription. Biophysical Journal, 121(9), 1583–1592. 10.1016/j.bpj.2022.03.026

Deb, A., Grewal, R. K., Roy, S., & Mitra, D. (2020). Residue interaction dynamics in *Vaucheria* aureochrome1 light-oxygen-voltage: Bridging theory and experiments. *Proteins: Structure*, Function, and Bioinformatics, 88(12), 1660–1674. 10.1002/prot.25984

Dmochowski, I. J., & Tang, X. (2007). Taking control of gene expression with light-activated oligonucleotides. BioTechniques, 43(2), 161–171. 10.2144/000112519

Dohno, C., Uno, S., Sakai, S., Oku, M., & Nakatani, K. (2009). The effect of linker length on binding affinity of a photoswitchable molecular glue for DNA. Bioorganic & Medicinal Chemistry, 17(6), 2536–2543. 10.1016/j.bmc.2009.01.053

Edwards, W. F., Young, D. D., & Deiters, A. (2009). Light-Activated Cre Recombinase as a Tool for the Spatial and Temporal Control of Gene Function in Mammalian Cells. ACS Chemical Biology, 4(6), 441–445. 10.1021/cb900041s

Erdős, G., & Dosztányi, Z. (2024). AIUPred: Combining energy estimation with deep learning for the enhanced prediction of protein disorder. Nucleic Acids Research, 52(W1), W176–W181. 10.1093/nar/gkae385

Gorostiza, P., & Isacoff, E. Y. (2008). Optical Switches for Remote and Noninvasive Control of Cell Signaling. Science, 322(5900), 395–399. 10.1126/science.1166022

Guerrero, L., Smart, O. S., Woolley, G. A., & Allemann, R. K. (2005). Photocontrol of DNA Binding Specificity of a Miniature Engrailed Homeodomain. Journal of the American Chemical Society, 127(44), 15624–15629. 10.1021/ja0550428

Halavaty, A. S., & Moffat, K. (2007). N- and C-Terminal Flanking Regions Modulate Light-Induced Signal Transduction in the LOV2 Domain of the Blue Light Sensor Phototropin 1 from *Avena sativa*. Biochemistry, 46(49), 14001–14009. 10.1021/bi701543e

Hart, J. E., Sullivan, S., Hermanowicz, P., Petersen, J., Diaz-Ramos, L. A., Hoey, D. J., Łabuz, J., & Christie, J. M. (2019). Engineering the phototropin photocycle improves photoreceptor performance and plant biomass production. Proceedings of the National Academy of Sciences, 116(25), 12550–12557. 10.1073/pnas.1902915116

Heintz, U., & Schlichting, I. (2016). Blue light-induced LOV domain dimerization enhances the affinity of Aureochrome 1a for its target DNA sequence. eLife, 5, e11860. 10.7554/eLife.11860

Herman, E., Sachse, M., Kroth, P. G., & Kottke, T. (2013). Blue-Light-Induced Unfolding of the Jα Helix Allows for the Dimerization of Aureochrome-LOV from the Diatom *Phaeodactylum tricornutum*. Biochemistry, 52(18), 3094–3101. 10.1021/bi400197u

Huysman, M. J. J., Fortunato, A. E., Matthijs, M., Costa, B. S., Vanderhaeghen, R., Van Den Daele, H., Sachse, M., Inzé, D., Bowler, C., Kroth, P. G., Wilhelm, C., Falciatore, A., Vyverman, W., & De Veylder, L. (2013). AUREOCHROME1a-Mediated Induction of the Diatom-Specific Cyclin *dsCYC2* Controls the Onset of Cell Division in Diatoms (*Phaeodactylum tricornutum*). The Plant Cell, 25(1), 215–228. 10.1105/tpc.112.106377

Im, S. H., Lepetit, B., Mosesso, N., Shrestha, S., Weiss, L., Nymark, M., Roellig, R., Wilhelm, C., Isono, E., & Kroth, P. G. (2024). Identification of promoter targets by Aureochrome 1a in the diatom *Phaeodactylum tricornutum*. Journal of Experimental Botany, 75(7), 1834–1851. 10.1093/jxb/erad478

Jolma, A., Yan, J., Whitington, T., Toivonen, J., Nitta, K. R., Rastas, P., Morgunova, E., Enge, M., Taipale, M., Wei, G., Palin, K., Vaquerizas, J. M., Vincentelli, R., Luscombe, N. M., Hughes, T. R., Lemaire, P., Ukkonen, E., Kivioja, T., & Taipale, J. (2013). DNA-Binding Specificities of Human Transcription Factors. Cell, 152(1–2), 327–339. 10.1016/j.cell.2012.12.009

Jumper, J., Evans, R., Pritzel, A., Green, T., Figurnov, M., Ronneberger, O., Tunyasuvunakool, K., Bates, R., Žídek, A., Potapenko, A., Bridgland, A., Meyer, C., Kohl, S. A. A., Ballard, A. J., Cowie, A., Romera-Paredes, B., Nikolov, S., Jain, R., Adler, J., … Hassabis, D. (2021). Highly accurate protein structure prediction with AlphaFold. Nature, 596(7873), 583–589. 10.1038/s41586-021-03819-2

Kalvaitis, M. E., Johnson, L. A., Mart, R. J., Rizkallah, P., & Allemann, R. K. (2019). A Noncanonical Chromophore Reveals Structural Rearrangements of the Light-Oxygen-Voltage Domain upon Photoactivation. Biochemistry, 58(22), 2608–2616. 10.1021/acs.biochem.9b00255

Khamaru, M., Deb, A., & Mitra, D. (2022). Basic Leucine Zippers: Aureochromes Versus the Rest. 10.1101/2022.05.19.492614

Khamaru, M., Nath, D., Mitra, D., & Roy, S. (2024). Assessing Combinatorial Diversity of Aureochrome Basic Leucine Zippers through Genome-Wide Screening. Cells Tissues Organs, 213(2), 133–146. 10.1159/000527593

Kobayashi, I., Nakajima, H., & Hisatomi, O. (2020). Molecular Mechanism of Light-Induced Conformational Switching of the LOV Domain in Aureochrome-1. Biochemistry, 59(28), 2592–2601. 10.1021/acs.biochem.0c00271

Kuravsky, M., Kelly, C., Redfield, C., & Shammas, S. L. (2024). The transition state for coupled folding and binding of a disordered DNA binding domain resembles the unbound state. Nucleic Acids Research, 52(19), 11822–11837. 10.1093/nar/gkae794

Laskowski, R. A., & Swindells, M. B. (2011). LigPlot+: Multiple Ligand–Protein Interaction Diagrams for Drug Discovery. Journal of Chemical Information and Modeling, 51(10), 2778–2786. 10.1021/ci200227u

Liang, X., Asanuma, H., & Komiyama, M. (2002). Photoregulation of DNA Triplex Formation by Azobenzene. Journal of the American Chemical Society, 124(9), 1877–1883. 10.1021/ja011988f

Lommer, M., Specht, M., Roy, A.-S., Kraemer, L., Andreson, R., Gutowska, M. A., Wolf, J., Bergner, S. V., Schilhabel, M. B., Klostermeier, U. C., Beiko, R. G., Rosenstiel, P., Hippler, M., & LaRoche, J. (2012). Genome and low-iron response of an oceanic diatom adapted to chronic iron limitation. Genome Biology, 13(7), R66. 10.1186/gb-2012-13-7-r66

Losi, A., & Gärtner, W. (2011). Old Chromophores, New Photoactivation Paradigms, Trendy Applications: Flavins in Blue Light-Sensing Photoreceptors^†^. Photochemistry and Photobiology, 87(3), 491–510. 10.1111/j.1751-1097.2011.00913.x

Martinez-Yamout, M. A., Nasir, I., Shnitkind, S., Ellis, J. P., Berlow, R. B., Kroon, G., Deniz, A. A., Dyson, H. J., & Wright, P. E. (2023). Glutamine-rich regions of the disordered CREB transactivation domain mediate dynamic intra- and intermolecular interactions. Proceedings of the National Academy of Sciences, 120(47), e2313835120. 10.1073/pnas.2313835120

Mayer, G., & Heckel, A. (2006). Biologically Active Molecules with a “Light Switch.” Angewandte Chemie International Edition, 45(30), 4900–4921. 10.1002/anie.200600387

Mészáros, B., Erdős, G., & Dosztányi, Z. (2018). IUPred2A: Context-dependent prediction of protein disorder as a function of redox state and protein binding. Nucleic Acids Research, 46(W1), W329–W337. 10.1093/nar/gky384

Mikat, V., & Heckel, A. (2007). Light-dependent RNA interference with nucleobase-caged siRNAs. RNA, 13(12), 2341–2347. 10.1261/rna.753407

Mitra, D., Yang, X., & Moffat, K. (2012). Crystal Structures of Aureochrome1 LOV Suggest New Design Strategies for Optogenetics. Structure, 20(4), 698–706. 10.1016/j.str.2012.02.016

Möglich, A., Ayers, R. A., & Moffat, K. (2009). Structure and Signaling Mechanism of Per-ARNT-Sim Domains. Structure, 17(10), 1282–1294. 10.1016/j.str.2009.08.011

Möglich, A., & Moffat, K. (2010). Engineered photoreceptors as novel optogenetic tools. Photochemical & Photobiological Sciences, 9(10), 1286–1300. 10.1039/c0pp00167h

Morgan, S.-A., Al-Abdul-Wahid, S., & Woolley, G. A. (2010). Structure-Based Design of a Photocontrolled DNA Binding Protein. Journal of Molecular Biology, 399(1), 94–112. 10.1016/j.jmb.2010.03.053

Papanatsiou, M., Petersen, J., Henderson, L., Wang, Y., Christie, J. M., & Blatt, M. R. (2019). Optogenetic manipulation of stomatal kinetics improves carbon assimilation, water use, and growth. Science, 363(6434), 1456–1459. 10.1126/science.aaw0046

Pettersen, E. F., Goddard, T. D., Huang, C. C., Couch, G. S., Greenblatt, D. M., Meng, E. C., & Ferrin, T. E. (2004). UCSF Chimera—A visualization system for exploratory research and analysis. Journal of Computational Chemistry, 25(13), 1605–1612. 10.1002/jcc.20084

Pinheiro, A. V., Baptista, P., & Lima, J. C. (2008). Light activation of transcription: Photocaging of nucleotides for control over RNA polymerization. Nucleic Acids Research, 36(14), e90–e90. 10.1093/nar/gkn415

Podust, L. M., Krezel, A. M., & Kim, Y. (2001). Crystal Structure of the CCAAT Box/Enhancer-binding Protein β Activating Transcription Factor-4 Basic Leucine Zipper Heterodimer in the Absence of DNA. Journal of Biological Chemistry, 276(1), 505–513. 10.1074/jbc.M005594200

Pomerantz, J. L., Sharp, P. A., & Pabo, C. O. (1995). Structure-Based Design of Transcription Factors. Science, 267(5194), 93–96. 10.1126/science.7809612

Rockwell, N. C., & Lagarias, J. C. (2010). A Brief History of Phytochromes. ChemPhysChem, 11(6), 1172–1180. 10.1002/cphc.200900894

Salomon, M., Christie, J. M., Knieb, E., Lempert, U., & Briggs, W. R. (2000). Photochemical and Mutational Analysis of the FMN-Binding Domains of the Plant Blue Light Receptor, Phototropin. Biochemistry, 39(31), 9401–9410. 10.1021/bi000585+

Sang, M., & Johnson, M. E. (2024). Oligomerization of transcription factors increases binding probability and residence time on DNA. Biophysical Journal, 123(3), 502a. 10.1016/j.bpj.2023.11.3034

Schellenberger Costa, B., Jungandreas, A., Jakob, T., Weisheit, W., Mittag, M., & Wilhelm, C. (2013). Blue light is essential for high light acclimation and photoprotection in the diatom Phaeodactylum tricornutum. Journal of Experimental Botany, 64(2), 483–493. 10.1093/jxb/ers340

Shestopalov, I. A., Sinha, S., & Chen, J. K. (2007). Light-controlled gene silencing in zebrafish embryos. Nature Chemical Biology, 3(10), 650–651. 10.1038/nchembio.2007.30

Shoemaker, B. A., Portman, J. J., & Wolynes, P. G. (2000). Speeding molecular recognition by using the folding funnel: The fly-casting mechanism. Proceedings of the National Academy of Sciences, 97(16), 8868–8873. 10.1073/pnas.160259697

Sreerama, N., & Woody, R. W. (2000). Estimation of Protein Secondary Structure from Circular Dichroism Spectra: Comparison of CONTIN, SELCON, and CDSSTR Methods with an Expanded Reference Set. Analytical Biochemistry, 287(2), 252–260. 10.1006/abio.2000.4880

Takahashi, F., Yamagata, D., Ishikawa, M., Fukamatsu, Y., Ogura, Y., Kasahara, M., Kiyosue, T., Kikuyama, M., Wada, M., & Kataoka, H. (2007). AUREOCHROME, a photoreceptor required for photomorphogenesis in stramenopiles. Proceedings of the National Academy of Sciences, 104(49), 19625–19630. 10.1073/pnas.0707692104

Taylor, B. L., & Zhulin, I. B. (1999). PAS Domains: Internal Sensors of Oxygen, Redox Potential, and Light. Microbiology and Molecular Biology Reviews, 63(2), 479–506. 10.1128/MMBR.63.2.479-506.1999

Tuszynska, I., Magnus, M., Jonak, K., Dawson, W., & Bujnicki, J. M. (2015). NPDock: A web server for protein–nucleic acid docking. Nucleic Acids Research, 43(W1), W425–W430. 10.1093/nar/gkv493

Waterhouse, A., Bertoni, M., Bienert, S., Studer, G., Tauriello, G., Gumienny, R., Heer, F. T., de Beer, T. A. P., Rempfer, C., Bordoli, L., Lepore, R., & Schwede, T. (2018). SWISS-MODEL: Homology modelling of protein structures and complexes. Nucleic Acids Research, 46(W1), W296–W303. 10.1093/nar/gky427

Williams, C. J., Headd, J. J., Moriarty, N. W., Prisant, M. G., Videau, L. L., Deis, L. N., Verma, V., Keedy, D. A., Hintze, B. J., Chen, V. B., Jain, S., Lewis, S. M., Arendall, W. B., Snoeyink, J., Adams, P. D., Lovell, S. C., Richardson, J. S., & Richardson, D. C. (2018). MolProbity: More and better reference data for improved all-atom structure validation. Protein Science, 27(1), 293–315. 10.1002/pro.3330

Wu, D., Potluri, N., Lu, J., Kim, Y., & Rastinejad, F. (2015). Structural integration in hypoxia-inducible factors. Nature, 524(7565), 303–308. 10.1038/nature14883

Yan, R., Xu, D., Yang, J., Walker, S., & Zhang, Y. (2013). A comparative assessment and analysis of 20 representative sequence alignment methods for protein structure prediction. Scientific Reports, 3(1), 2619. 10.1038/srep02619

Zhang, H., Xiong, X., Guo, K. et al. (2024) A rapid aureochrome opto-switch enables diatom acclimation to dynamic light. Nat Commun 15, 5578. 10.1038/s41467-024-49991-7

Zoltowski, B. D., Vaccaro, B., & Crane, B. R. (2009). Mechanism-based tuning of a LOV domain photoreceptor. Nature Chemical Biology, 5(11), 827–834. 10.1038/nchembio.210

