## Supplementary Information for "Exploring the Modularity of *Thalassiosira oceanica* Aureochrome1 Photoreceptor-Transcription Factor: Characterization and Component Engineering"

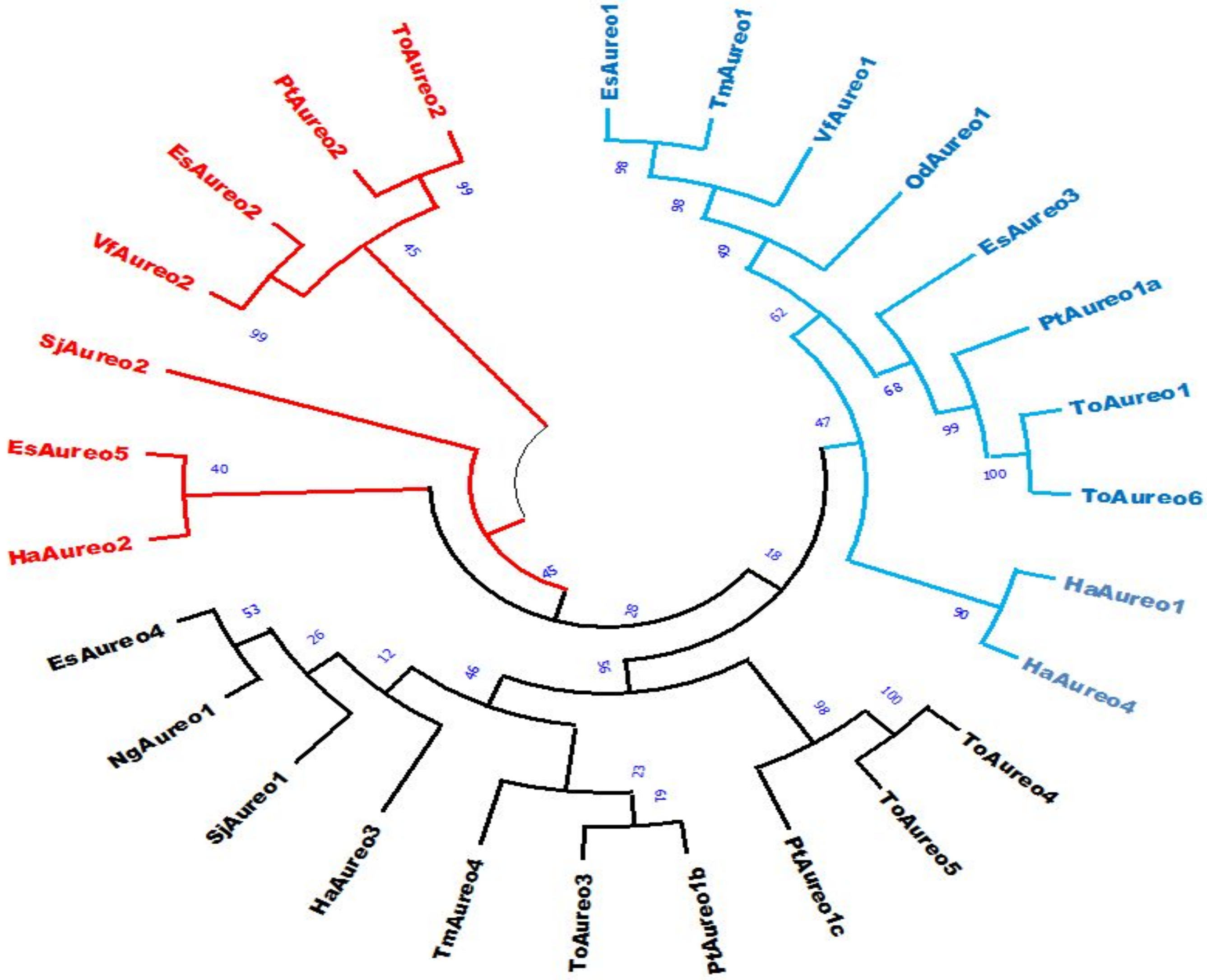


**Figure S1:** Phylogenetic tree of Aureochromes showing distinct clustering of Aureo2 from the rest. Other Aureos, including Aureo1 are clubbed together.

**N-(X)7-R/K**

```
EsAureo1    GRGPHSGAGATAAPRRQRHKVSSKDL---------------------TEEQRIERRERNREHAKRSRVRKKFLLDSLQRSVDAIQAENESLKGSIVGSLGERGRELVAKCMPADTGG---
TmAureo1    NRGVHAGIGGGGGG-------SSAPL---------------------TEAQKAERREKNREHAKRSRVRKKFLLESLQRSVDTLQNENASLKASIREYMGETGDELIAKCVPEGED----
VfAureo1    IYESQGNASRGKSLRTKSSGSISSEL---------------------TEAQKVERRERNREHAKRSRVRKKFLLESLQQSVNELNHENNCLKESIREHLGPRGDSLIAQCSP--EAD---
SjAureo2    -------------------------M---------------------TEQQRLDRRERNREHAKRSRVRKKFLLESLQKSVTALQEENEKLRGAIRANLGADEAKELLAQTES-------
EsAureo3    -------GGKRGSVSGKRPRQGGRNM---------------------TEQQRLDRRERNREHAKRSRVRKKFLLESLQKSVTSLQEENEKLRGAIRSNLGPEEAKELLAETETP------
ToAureo1    ATSNAGKKRAIGTSAATATKPPVEKK---------------------SMDR----RERNREHAKRSRIRKKFLLESLQQSVSLLKEENGKLKNAIRTHLGEKEAEALLNKQALADAASSG
ToAureo6    ATSNAGKKRAIGTSAATATKPPVEKK---------------------SMDRSQVVKERNREHAKRSRIRKKFLLESLQQSVSLLKEENGKLKNAIRTHLGEKEAEALLNKQALADAASSG
PtAureo1a   ------KKKSSSSSTSGSGSRSDRKS---------------------MEQQKVERRERNREHAKRSRIRKKFLLESLQQSVSLLKEENEKLKTSIRSHLGDEKADTLIDSANNNKTDVDG
OdAureo1    ------EGDSDDGKRRKRSRTLPRNM---------------------TEEQKIERRERNREHAKRSRVRKKFLLESLQHSVRALEEENEKLRNAIRENLQGEAEQLLTRCSCGG------
HaAureo1    QRKRPKVPQATISETAIFKGFHGVEP---------------------AKIAEKIRKQRNKEHAKRSRVRKKFLVDSLQQSIDLLEKENQKLRSCLSANLGPEAQSIIHAGVTECSSLIAH
HaAureo4    -----------------------------------------------------MRKERNKEHAKRSRTRKKFLLDSFQYALDVLTKENEALRKNITASLGEEEAEREFAAFAELHRANAA
SjAureo1    ----------------------MSEQ------------------------QKLERRERNREHAKRSRIRKKFMLECLQEQLLAMRKQNMALRQVVKEHMPNEAATVFEKCVHEKALALSG
EsAureo4    DARR-----------------SMSEQ------------------------QKLERRERNREHAKRSRIRKKFMLECLQEQLLAMRKQNLALRQVVKEHMPDEASTVFEKCINEKALVLSG
TmAureo4    ERKRKASGAEGHVNLDKLPISELTEE------------------------QKIERRERNREHAKRSRVRKKFMLDCLQDQLLALRKENMNLRQVVKEHMPEVATQIFEKCVSEKALVLSA
NgAureo1    EGEGGDGRGARRRKKAKLTLKPLTEA------------------------QRVERRERNREHAKRSRMRKKFMLESLQAQMLALRKENLRLRQLVATKLPDKADTILRGCSSIKTQNLLS
HaAureo3    NFNDTSQSAASAAAAQQGTKRNLTEE------------------------QRLERRERNREHAKRSRIRKRFMLESLQEQFMGLQRENMALRQIIKDKIPARADSIFEKCTDKISDLIST
ToAureo4    RNSARKPAPSTERGLSTRPAASLTES------------------------QKTERRERNREHAKRSRLRKKFLLESLQEQIDGLQGEISTLKSIIKRELPDRAAALLKDIGDSDEADSAA
ToAureo5    RNSARKPAPSTERGLSTRPAASLTES------------------------QKTERRERNREHAKRSRLRKKFLLESLQEQIDGLQGEISTLKSIIKRELPDRAAALLKDIGDSDEADSAA
PtAureo1c   -----------------TGNKDLTEQ------------------------QKLERRERNREHAKRSRLRKKFLLESLQEQIHGLEEQLDGLKSAIKKELPQQAEQIITRICG--------
PtAureo1b   PATAQSATRTSGVPGQFNMWGPGTGG--------------------MSEQAVAERRQRNREHAKRSRVRKKFMLESLQEQVREMQKQNQNLRMLVQEHIPEHAMKIIAECCTSSPLFEEM
ToAureo3    QQTLQNFGTNTSVLQAALGGGPLHVTNVQYGVNPLQQQHVVTKHLNPEESTQTERRLRNREHAKRSRVRKKFMLESLQQQVKGLQDENGTLRLLVQKHIPKHALKIIDECCSKSILFRTA
PtAureo2    ARAIQNQG-----------RGRKKSQ-----------------------AQIDRRRERNRIIARRTRLRKKFFFESLQKEIMDLQRENLVLKELVKVNISGEEGKKILEGCNAAENLPSS
ToAureo2    MAAQINKAHPMPPAPPTDSRSGEKSQ-----------------------AQVDRRRERNRILARRTRLRKKFFFESLQKDVADLQRENAALKSIVRSRLKPGDARSVLDQCDAHKKMPGD
VfAureo2    SQECLNTAYKSDATTATTRRRAQSRE-----------------------EQQQKRRERNKVLARKTRLRKKFLFQGLRNQVMSLYQENLRLKEIVKNHCGINSKKILDDNCLGDMPDIVA
EsAureo2    ALGGVGSTFRKTPAGGVTRRRSSSKE-----------------------EQAKKRRERNRVLARRTRLRKKFFFQSLQQQVARLQRENERLKGIVTTRCPDSVGEILMS-CRSKMPSMVA
HaAureo2    LLQQQQQQPSQSDEPGRKKGNKLTPE-----------------------EQVERRRERNRILARKTRLRKKLFFEALQRRVTNLKTENELLRGVAQRRLGDADRRAALGGLHSELPKVVT
EsAureo5    --------MIMRGIPDRAG---------------------------------AIIRNCAKDNTNADVVAKQDVIQGMRPGMSVGSSSSSWTARGGTRTRPKPAGGGGGGGGSGKGRYVPS
                                                                :    :   :.      *:  ..:  ::   .       .
```

**Basic Region** **Leucine Zipper Region**

**Figure S2:** Multiple sequence alignment of Aureochrome bZIPs showing highly conserved basic region. The unique histidine substitution is also prevalent in all Aureos except Aureo2.

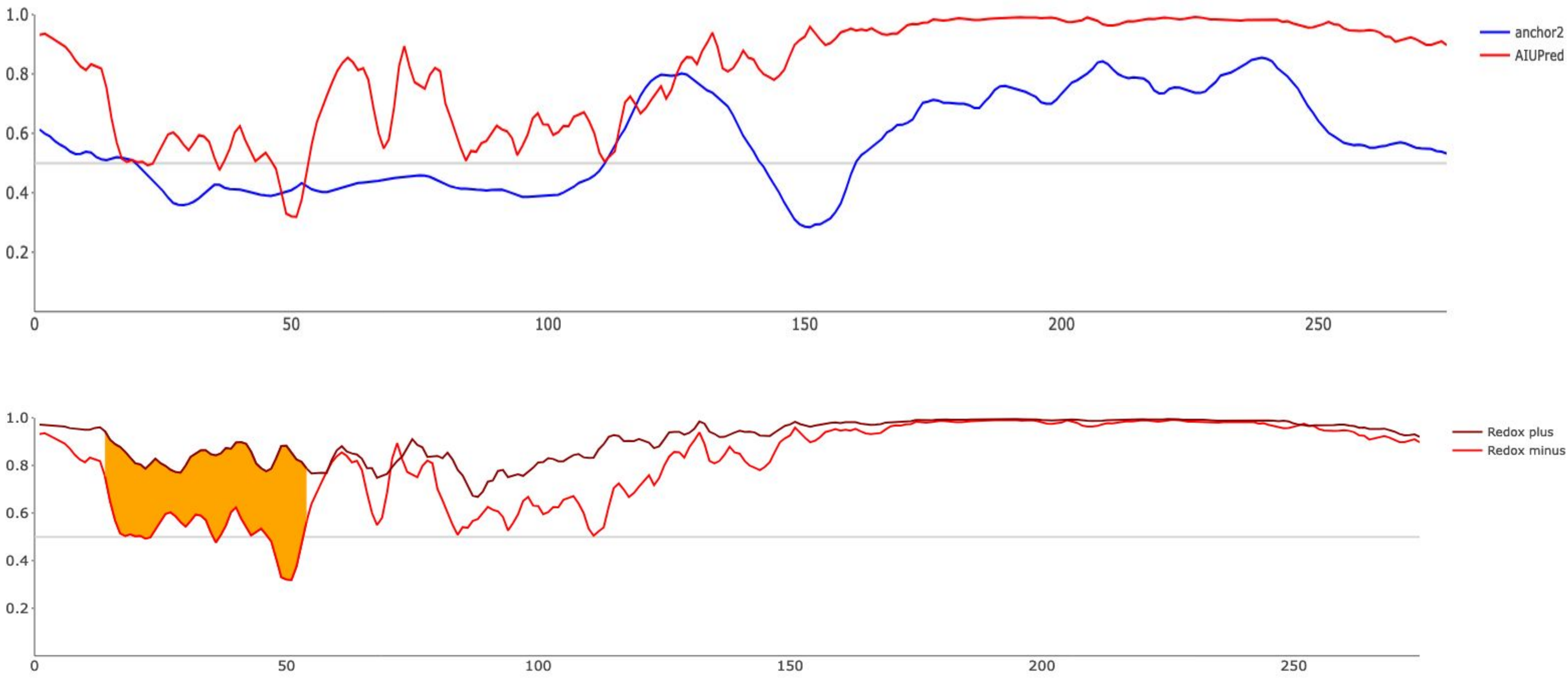


**Figure S3:** AIUPRED results of *To*Aureo1 N-terminal extension (NTE) reveal largely disordered nature. Disordered binding regions are noticed in between 111 to 141 preceding the glutamine-rich sequence and again between 160th residue till the end of NTE (*top*). 18-53rd residue stretch further shows redox sensitivity (*bottom*).

```
EsAureo1     --SLVTANN--------------------KQATKILDDPDYSLVKALQTAQQNFVITDAS
TmAureo1     --DLVTSNP--------------------SRATRVLDDPDYSLVKALQLAQQNFVITDAS
VfAureo1     --TLLTDNP--------------------SKANRILEDPDYSLVKALQMAQQNFVITDAS
SjAureo2     -SSLIASNP--------------------GDATKVLDDPDYSLVKALQTAQQNFVITDPT
EsAureo3     -TTLISSRP--------------------GDATKVLDDPDYSLVKALQTAQQNFVITDPT
ToAureo1     GGGLIANNK--------------------SDANRVLDDPDFSFIKALQTAQQNFVVTDPS
ToAureo6     GGGLIANNK--------------------SDANRVLDDPDFSFIKALQTAQQNFVVTDPS
PtAureo1a    ---LLASSQ--------------------GIANKVLDDPDFSFIKALQTAQQNFVVTDPS
OdAureo1     -PSVIASDP--------------------NTATRTLDDPDYSLVKALQTAQQNFVISDPS
HaAureo1     DFAS----------------------------AKVLEDPDFSLVTALQSAQKSFVITDPA
HaAureo4     GAAVDEDDDAAVQQPALFGPDGQAASAEGGGGGTGLEAEDAGLLRSLQKARFSFVISNPA
SjAureo1     KGTGSAQAIQSDD------------EDESKEPNCLLLEPDFQLMQALMESQQNFTISDPS
EsAureo4     KSTESAQPILSDD------------EEETKEPNCLLLEPDFQLMQALMTSQQNFTISDPS
TmAureo4     DTGEAAAGGHAAV------------AFTSK-PDCLLVEPDFQLISALMDSQQNFTISNPH
NgAureo1     S--------------------------VELGHHRALADHDGRLVSALQFAQQNFTVSDPS
HaAureo3     P-------------------------VKDYQALRFLIEPDFRLVESLTTSQQNFTVSDPS
ToAureo4     PNKGSFTPLP---------------MPSGFGPVKTLMEPDYRLMSALSGSQQNFAISDPS
ToAureo5     PNKGSFTPLP---------------MPSGFGPVKTLMEPDYRLMSALSGSQQNFAISDPS
PtAureo1c    -DKEKFTPLP---------------MPSGFGPVKTLMEPDFRLMSALSGSQQNFAISDPT
PtAureo1b    DGIDQTKGAN-------------------------LERADFSLMQSLTMGQQCFVLSDPK
ToAureo3     DGANAGKKDDK------------------DAKETPLLRSDFSLIESLTSGQQNFVLSDPR
PtAureo2     VLEACGEEND-------------------------MDSQDFNLVRSIQSSQHSFMITDPS
ToAureo2     VRSSQFKENDGTNLTGA--------APDGGSSVKKLDREDFSLIQSIQNSQQCFIISDPS
VfAureo2     TCGQEATALIQ--------------------------RSDFTLMKALYSSQPSFCVTNPL
EsAureo2     DCAGQATAVLD--------------------------QSGFLLVKALQSSQPSFCVTDPQ
HaAureo2     ENMGQATEVIK--------------------------KTDFAMMKLLTTAQKSFVITDPS
EsAureo5     --------------------------------ALKLMSPCYQTVKALADSQANFVLTNPN
                                                        :  :  .:  * :::.
```

Heterogeneous Linker Region Aureochromes

**Figure S4:** Multiple sequence alignment of Aureochrome bZIPs showing poorly conserved linker region connecting bZIP and linker.

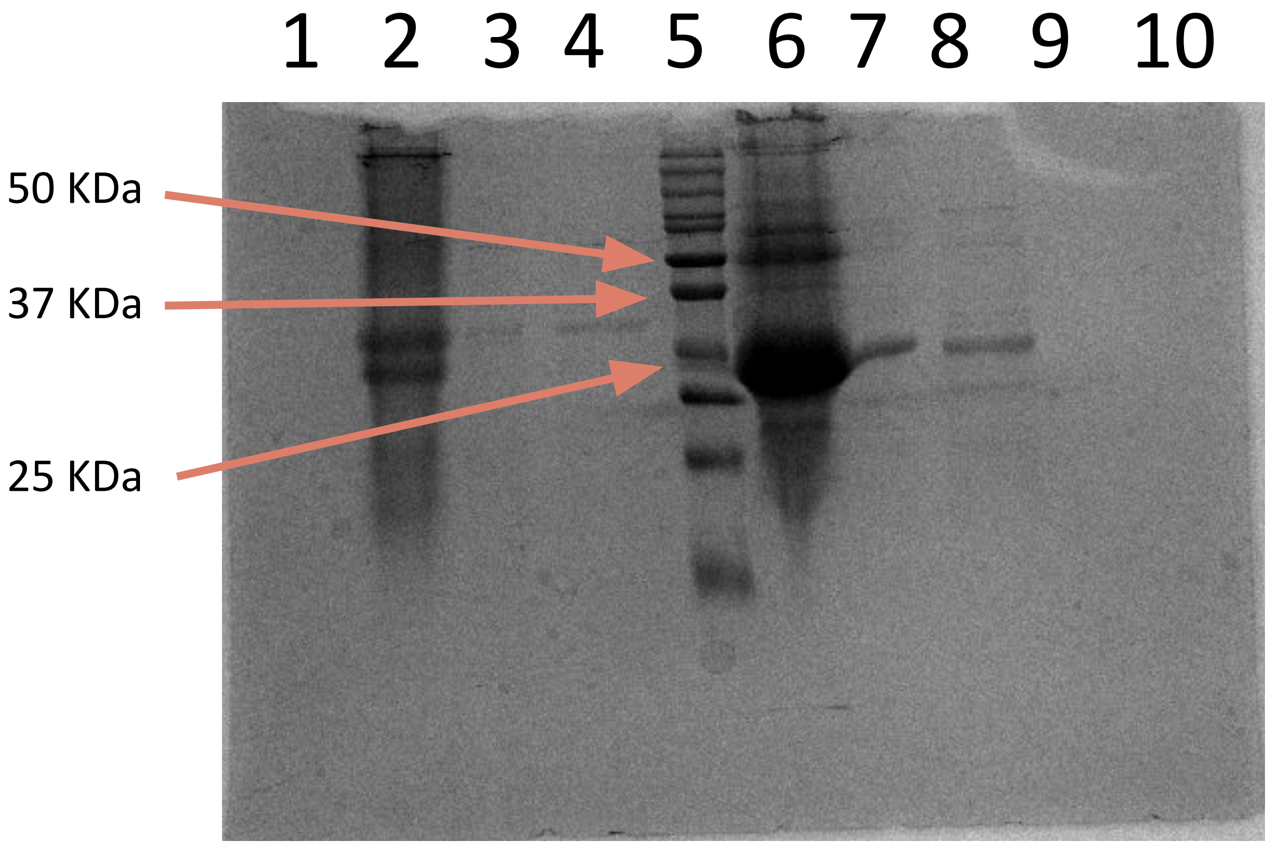


**Figure S5:** SDS-PAGE analysis of *To*AubZL and *AsTo*bZL. Lane 1, 9 and 10 - blank; lane 2 - *To*AubZL Ni-NTA fraction; lane 3 - *To*AubZL gel filtration fraction; lane 4 - *To*AubZL Heparin fraction; lane 5 - Protein molecular weight marker; lane 6 - *AsTo*bZL Ni-NTA fraction; lane 7 - *AsTo*bZL gel filtration fraction and lane 8 - *AsTo*bZL Heparin fraction.

### *To*AubZL (A)

| Secondary Structures (%) | Selcon 3 | Contin LL |
|---|---|---|
| Helix 1 | 1.6 | 1.1 |
| Helix 2 | 2.7 | 7.6 |
| Strand 1 | 19.2 | 22.8 |
| Strand 2 | 11.4 | 12.1 |
| Turns | 18.6 | 22.2 |
| Unordered | 31.2 | 34.1 |
| Total | 84.6 | 99.9 |

### *AsTo*bZL (B)

| Secondary Structures (%) | Selcon 3 | Contin LL |
|---|---|---|
| Helix 1 | 1.0 | 2.7 |
| Helix 2 | 2.3 | 5.9 |
| Strand 1 | 20.5 | 21.2 |
| Strand 2 | 12.9 | 10.4 |
| Turns | 24.0 | 17.7 |
| Unordered | 38.8 | 42.1 |
| Total | 99.4 | 100 |

**Table S1:** CD spectroscopy data of *To*AubZL and *AsTo*bZL analyzed by different methods showing similar content of secondary structural elements.

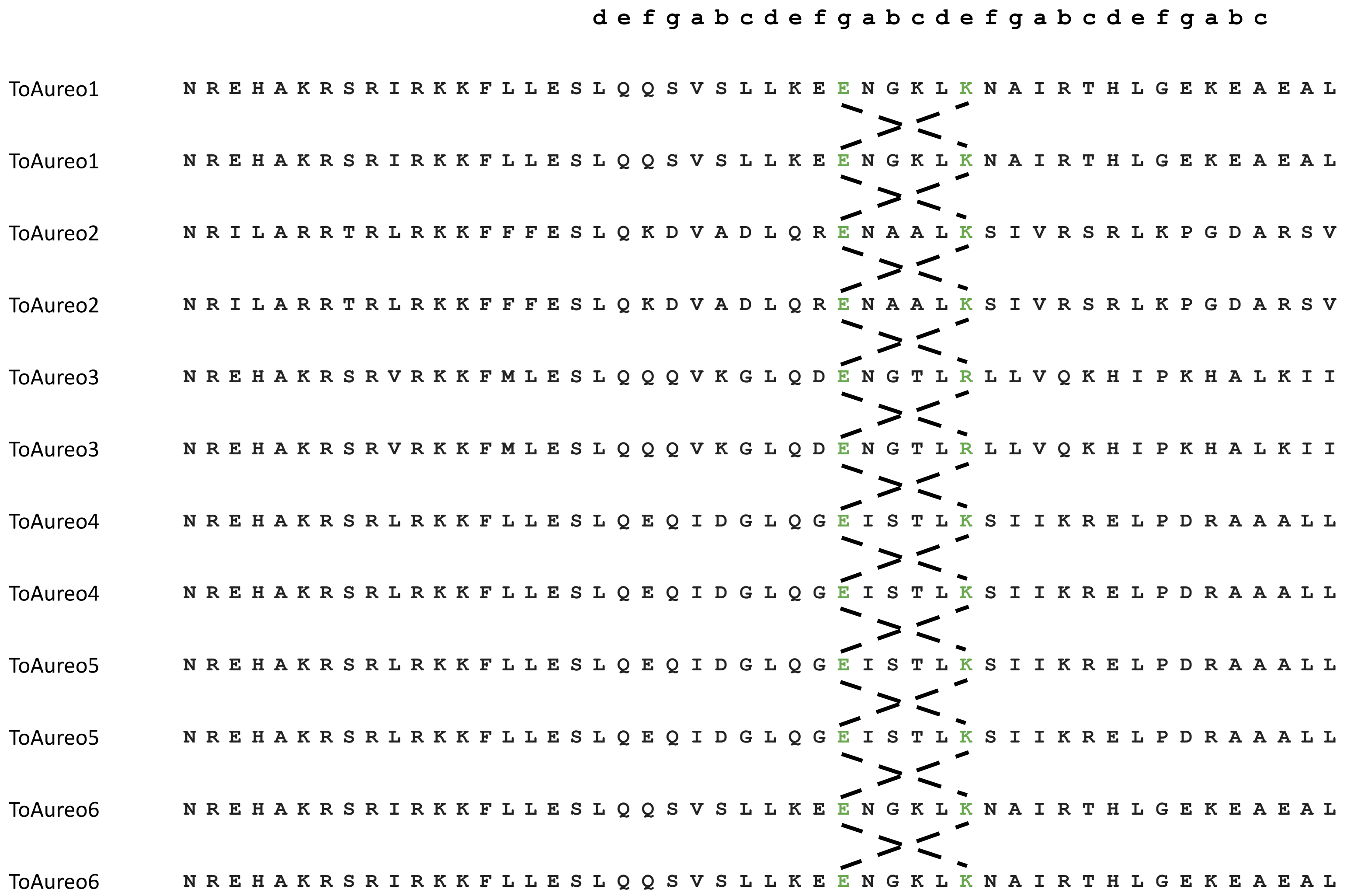


**Figure S6:** Compatibility analysis of *To*Aureo homo/heterodimerization shows high possibility of formation of homodimers in all combinations.
